# *In Vivo* CRISPR Screens Identify E3 Ligase Cop1 as a Modulator of Macrophage Infiltration and Cancer Immunotherapy Target

**DOI:** 10.1101/2020.12.09.418012

**Authors:** Xiaoqing Wang, Collin Tokheim, Binbin Wang, Shengqing Stan Gu, Qin Tang, Yihao Li, Nicole Traugh, Yi Zhang, Ziyi Li, Boning Zhang, Jingxin Fu, Tengfei Xiao, Wei Li, Clifford A. Meyer, Jun Chu, Peng Jiang, Paloma Cejas, Klothilda Lim, Henry Long, Myles Brown, X. Shirley Liu

## Abstract

Despite remarkable clinical efficacies of immune checkpoint blockade (ICB) in cancer treatment, ICB benefits in triple-negative breast cancer (TNBC) remain limited. Through pooled *in vivo* CRISPR knockout (KO) screens in syngeneic TNBC mouse models, we found that inhibition of the E3 ubiquitin ligase *Cop1* in cancer cells decreases the secretion of macrophage-associated chemokines, reduces tumor macrophage infiltration, and shows synergy in anti-tumor immunity with ICB. Transcriptomics, epigenomics, and proteomics analyses revealed *Cop1* functions through proteasomal degradation of the *C/ebpδ* protein. *Cop1* substrate *Trib2* functions as a scaffold linking *Cop1* and *C/ebpδ*, which leads to polyubiquitination of *C/ebpδ*. *Cop1* inhibition stabilizes *C/ebpδ* to suppress the expression of macrophage chemoattractant genes. Our integrated approach implicates *Cop1* as a target for improving cancer immunotherapy efficacy by regulating chemokine secretion and macrophage levels in the TNBC tumor microenvironment.

**Highlights:**

1. *Large-scale in vivo CRISPR screens identify new immune targets regulating the tumor microenvironment*
2. *Cop1 knockout in cancer cells enhances anti-tumor immunity*
3. *Cop1 modulates chemokine secretion and macrophage infiltration into tumors*
4. *Cop1 targets C/ebp*δ *degradation via Trib2 and influences ICB response*

## INTRODUCTION

Breast cancer is one of the leading causes of cancer-associated morbidity and mortality in the United States (Fallahpour et al., 2017; Waks and Winer, 2019). Triple-negative breast cancer (TNBC) constitutes 15% of breast cancer and has the worst prognosis among the molecular subtypes, motivating research efforts to find new treatment options in TNBC (Bianchini et al., 2016). Immune checkpoint blockade (ICB) has shown remarkable clinical benefits to skin, lung and colorectal cancer patients (Halle et al., 2017), raising the possibility of effective ICB treatment of breast cancer. In 2019, the FDA approved the first ICB therapy for the treatment of metastatic TNBC. Atezolizumab, an anti-PD-L1 monoclonal antibody, was approved in combination with nab-paclitaxel (nanoparticle albumin-bound paclitaxel) based on prolonged progression-free survival (Schmid et al., 2018). While this advance demonstrates the promise of ICB in breast cancer treatment, the benefits were limited to a small subset of patients. A recent clinical trial reported that pembrolizumab, an anti-PD-1 monoclonal antibody, had an objective response rate of just 18% in PD-L1-expressing advanced TNBC (Nanda et al., 2016). This underscores the need for finding new immune targets to enhance ICB response and improve outcomes in TNBC.

The immune system is thought to influence the progression of most cancer types (Coussens et al., 2013; Hanahan and Weinberg, 2011), but the molecular mechanisms governing tumor immunity and the tumor microenvironment (TME) are not fully understood. Some cell types in the TME are proposed to promote tumor growth and metastasis, such as myeloid-derived suppressor cells (MDSC) (Grivennikov et al., 2010), fibroblasts (Landskron et al., 2014), and tumor-associated macrophages (TAMs) (Su et al., 2018). Among them, TAMs are a major player and are thought to promote angiogenesis, cancer cell local invasion, and intravasation at primary tumor sites. At metastatic sites, TAMs can also facilitate cancer cell extravasation and block CD8^+^ T cell recruitment and functions (Cassetta and Pollard, 2018; Peranzoni et al., 2018). In patients, TAM infiltration is strongly associated with poor clinical outcomes in numerous cancer types, including breast cancer (Cassetta et al., 2019; Zhang et al., 2016). In syngeneic mouse models, classical monocytes (Mouse CD11b^+^Ly6C^+^) are recruited to tumor and differentiate into TAMs as tumors progress (Ginhoux and Jung, 2014). This process often depends on macrophage chemoattractants secreted from cancer cells or activated macrophages, such as CCL2 (Nielsen and Schmid, 2017), CCL4 (Li et al., 2018), CCL5 (Walens et al., 2019), CXCL1 (Wang et al., 2017), and CXCL5 (Zhao et al., 2017). Accordingly, a monoclonal antibody was developed to inhibit CCL2 signaling pathway, which indeed attenuates TAM infiltration, suppresses tumor growth, and improves survival (Qian et al., 2011). However, pharmacological inhibition of chemokines are associated with chemokine over-expression due to homeostatic feedback, yielding adverse effects (Lim et al., 2016). These findings motivated us to discover novel targets to reprogram the TME for cancer treatment.

Functional genomic screening using CRISPR-Cas9 has shown promise as a robust and unbiased approach to discover novel cancer targets. It has also been adopted to find novel modulators of tumor immunity, discover novel immuno-oncology targets, and dissect their mechanisms. For instance, genome-wide CRISPR screens in human pancreatic cancer cells *in vitro* have shown that *CMTM6* loss robustly inhibited *PD-L1* expression and enhanced anti-tumor immunity (Burr et al., 2017). CRISPR screens in cancer cells co-cultured with T cells identified *Pbrm1* loss as increasing B16F10 melanoma cell sensitivity to effector T cells (Pan et al., 2018), and other genes allowing cancer cells to escape the immune system (Lawson et al., 2020). Pooled *in vivo* CRISPR screens in the murine melanoma models revealed that loss of *Ptpn2* (Manguso et al., 2017) and *Adar1* (Ishizuka et al., 2019) could enhance tumor sensitivity to immunotherapy. While *in vivo* CRISPR screens can capture the broad landscape of tumor immunity, it can only screen a few hundred genes at a time, which has limited their application to a small number of tumor models. This encouraged us to test more genes in different syngeneic models using *in vivo* CRISPR screens, with the intention of identifying new regulators of tumor immunity.

In this study, we constructed and tested a custom murine CRISPR Knockout library *in vitro*, then conducted *in vivo* CRISPR screens in two murine TNBC models under different levels of host immunity. The *in vivo* screens identified genes involved in regulating an effective immune response against cancer cells in TNBC. We validated the effect of each identified target gene on tumor immune-cell infiltration and anti-PD-1 response *in vivo*. To further characterize the function and mechanisms of potential therapeutic targets, we performed RNA-seq, ATAC-seq, and proteomics analysis, and used this data to further narrow the candidates. Observations on clinical tumor immune infiltration and patient survival across many cancer types strongly support the clinical relevance of the identified candidate therapeutic target genes.

## RESULTS

### Large-scale *in vivo* CRISPR Screens Identify Regulators of Immune Evasion

To systematically discover novel gene targets in cancer cells that when disabled enhance anti-tumor immunity, we first constructed a murine CRISPR-Cas9 knockout (MusCK) library. This library includes 5 sgRNAs for each of over 4,500 genes implicated in tumor initiation, progression, and immune modulation (Figure S1A; Table S1) (see STAR Methods for further details). To validate the MusCK library, we introduced it by lentiviral transduction into the mouse TNBC cell line 4T1 *in vitro* (Figure S1B) (see STAR Methods for further details). 4T1 cells closely resemble human TNBC cells (Figures S1C and S1D), are transplantable into the syngeneic BALB/c background mice, and have been extensively used in tumor immunology studies (Kim et al., 2014; Sagiv-Barfi et al., 2015). We compared sgRNA abundance distribution in freshly infected 4T1 cells to the sgRNA abundance distribution in 4T1 cells cultured 10 passages after infection. Supporting the reliability of the MusCK library, cells harboring sgRNAs targeting known oncogenes and tumor suppressor genes were significantly depleted and enriched, respectively (Figure S1E and S1F; Table S2).

With the MusCK library validated, we next conducted *in vivo* CRISPR screens in 4T1 cells in syngeneic BALB/c mice. To this end, we first artificially expressed membrane-bound ovalbumin (mOva) in 4T1 cells, an approach widely used to enhance cellular immune responses in syngeneic tumor models. As expected, 4T1 tumors overexpressing mOva had increased lymphocyte infiltration and slower tumor growth (Figures S1G-S1J) (see STAR Methods for further details). We then transduced the lentiviral MusCK library into mOva-expressing 4T1 cells and implanted infected cells into the mammary fat pads of BALB/c Foxn1^nu/nu^ hosts (nude), BALB/c hosts, and BALB/c hosts vaccinated with ovalbumin prior to transplantation (Figure 1A; Figures S1K-S1L). We used 12 mice per arm and injected enough cells per mouse to achieve ∼200-fold coverage for all the sgRNAs in the MusCK library. Sixteen days post transplantation we harvested the engrafted cancer cells for analysis, and observed significantly different tumor growth in different hosts. While the T-cell deficient BALB/c Foxn1^nu/nu^ hosts had the biggest tumors, the immune-competent hosts pre-vaccinated with ovalbumin had the smallest tumors (Figure 1B; Figure S1M).

**Figure 1.**
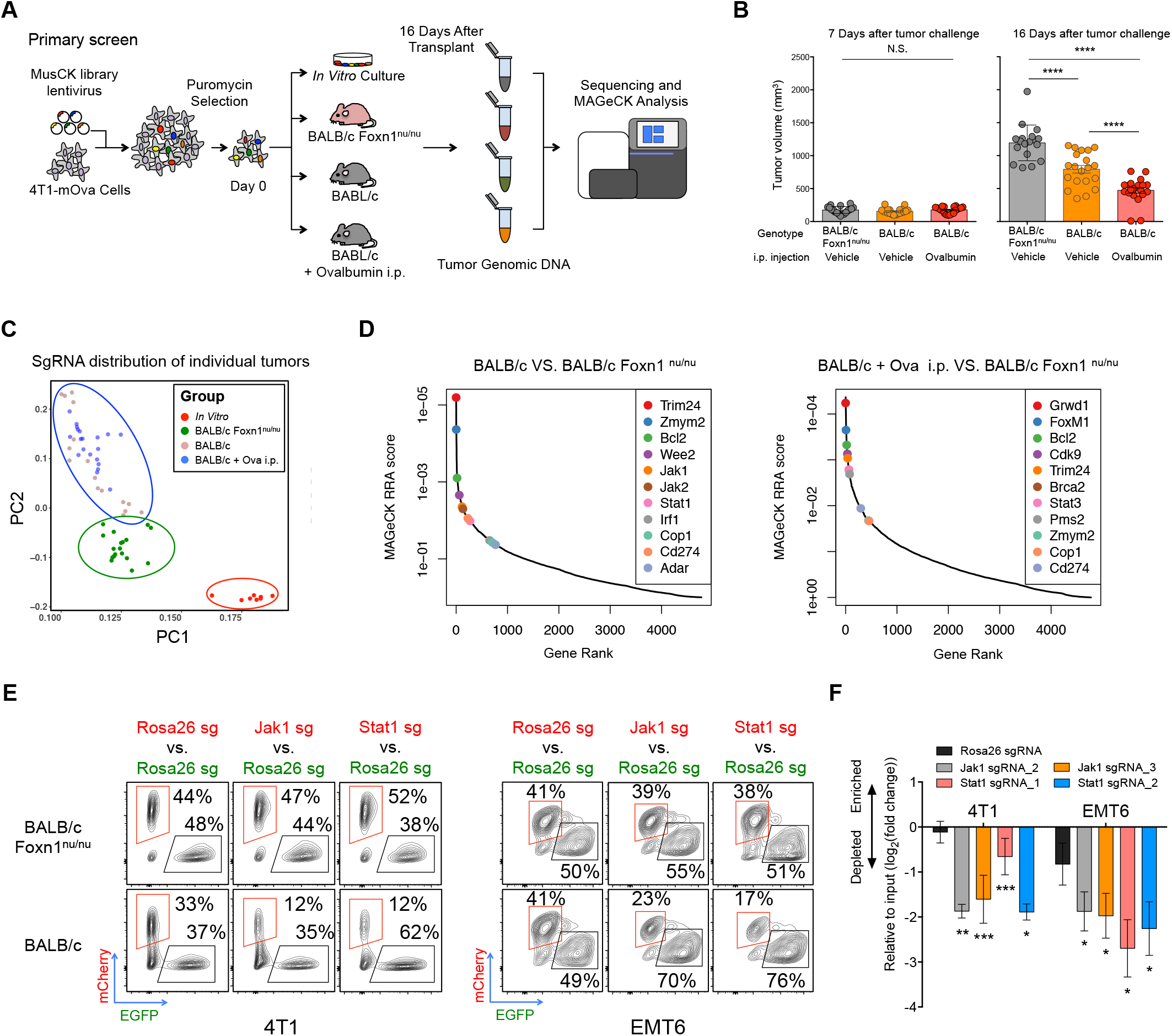
*In Vivo* Screens with the MusCK Library Uncovered Classic and Novel Regulators of Immune Evasion. (A) Workflow of using MusCK *in vivo* screens to identify the potential targets for immune evasion. i.p. = intraperitoneal. (B) Tumor volume measured at 7 and 16 days post implantation in the MusCK screens. Data are shown as mean ± SEM, n = 10-12 mice per group, * * * * P < 0.0001, by one-way ANOVA with Benjamini-Hochberg post-test multiple comparison. (C) Principal component analysis of sgRNA abundance in each condition of the MusCK screens. (D) Top depleted genes in immunocompetent versus immunodeficient (nude) hosts in the MusCK screens. (E) Flow cytometry analysis of Jak1 (or Stat1) KO cells versus control Rosa26 KO mouse breast cancer cells in the resulting 4T1 and EMT6 tumors. (F) Quantification of the relative percentages calculated from flow cytometry analysis. Data are shown as mean ± SEM, n = 4-6 mice per group, *p < 0.05, **p < 0.01, ***p < 0.001, by one-way ANOVA with Benjamini-Hochberg post-test multiple comparison.

To analyze the CRISPR screen results, we examined the sgRNA abundance distribution in the resulting 4T1 tumors grown *in vivo*. Reflective of different selection pressure in the hosts, principal component analysis showed that CRISPR screen samples separated first by *in vitro* versus *in vivo* conditions, then by nude mice versus immunocompetent mice (Figure 1C). Samples from the same condition cluster together, indicating similar library representations in biological replicates (Figure 1C). Inspection of the sgRNAs depleted from tumors in wild-type immunocompetent hosts compared to nude immunodeficient mice revealed key genes promoting immune evasion in 4T1 cancer cells (Figure 1D; Table S3) (see STAR Methods for further details). As positive controls, the sgRNAs targeting *Cd274* (*Pd-l1*) were depleted in tumors engrafted in wild-type mice, consistent with the known function of *Cd274* in immune suppression (Dong et al., 1999; Freeman et al., 2000). In addition, key components of DNA repair pathways, such as *Brca2* and *Pms2*, were significantly negatively selected in wild-type mice (Figure 1D). This is also consistent with previous reports that cancer cells with greater genome instability or mutation burden were at risk of elimination by T-cell mediated killing (Mandal et al., 2019; Pearlman et al., 2017).

Interestingly, key components of the IFNγ pathway (*Jak1, Jak2, Stat1 and Irf1*) were significantly depleted in wild-type mice, but not nude mice (Figure 1D), suggesting that defects in the IFNγ pathway in cancer cells could suppress immune evasion. IFNγ is a cytokine secreted by tumor-infiltrating lymphocytes to elicit anti-tumor immune response (Alshaker and Matalka, 2011). This result is opposite to findings from CRISPR-mediated genetic KO screens in the murine B16F10 melanoma model (Manguso et al., 2017), but is consistent with the role of IFNγ in promoting tumor immune evasion in multiple cancer types (Beatty and Paterson, 2000; Benci et al., 2016, 2019). There have been reports that the duration of IFNγ signaling contributes to differential tumor response to ICB (Minn, 2015). RNA-seq analysis revealed that IFNγ signaling was active in 4T1 tumors *in vivo* but not in 4T1 cells *in vitro* (Figure S1N). Thus, it is possible that prolonged IFNγ signaling in the tumors has immunosuppressive function, which explains why KO of IFNγ pathway genes enhances immune-mediated killing of TNBC cells. To confirm our findings, we conducted a competition assay to assess the *in vivo* growth of TNBC cells deficient in IFNγ signaling (Figure S1O). Western blotting confirmed the protein abundance of *Jak1* or *Stat1* KO in TNBC cells (Figure S1P). Then, we mixed cancer cells (1:1 ratio, mCherry:eGFP) with *Jak1* (or *Stat1*) KO and control *Rosa26* KO (see STAR Methods for further details), and implanted the cell mixture into nude and wild-type mice. Flow cytometry analysis in the resulting tumors showed that the relative proportion of *Jak1* or *Stat1* KO cancer cells became consistently and significantly lower than those of control cells (Figures 1E and 1F) especially in the wild-type mice. The same result was observed in another TNBC syngeneic model EMT6 (Figures 1E and 1F), which not only supports the reliability of *in vivo* screens using our MusCK library, but also confirms the role of IFNγ signaling in suppressing an anti-tumor immune response in TNBC.

### Loss of *Cop1* Sensitizes Cancers to Immunotherapy

Achieving adequate statistical significance for discovery in large-scale CRISPR screens requires behavioral consistency of multiple sgRNAs, each with sufficient cell coverage, for each target gene. To improve the robustness of our *in vivo* CRISPR screens, we constructed a second library (MusCK 2.0) focused on 79 candidate genes identified in the primary screen, with 8 sgRNAs per gene (see STAR Methods for further details). We then conducted a validation screen using the MusCK 2.0 library in 4T1-mOva cells implanted into (1) BALB/c Foxn1^nu/nu^ nude hosts; (2) wild-type BALB/c hosts; (3) wild-type BALB/c hosts with ovalbumin pre-vaccination; and (4) wild-type BALB/c hosts with both ovalbumin pre-vaccination and monoclonal anti-PD-1 treatment (Figure S2A). The additional fourth group facilitates the discovery of factors that affect antigen-specific T-cell immunity through the PD-1/PD-L1 axis. As expected, we observed statistically significant and progressively lower tumor volumes in groups (1) through (4) at 16 days after cancer cell implantation (Figure 2A; Figures S2A-S2C). We also observed progressively higher T-cell infiltration (detected by TCRβ^+^) relative to the total tumor immune infiltrates (marked by Cd45.2^+^) (Figure 2B; Figures S2D-S2F) in these four groups. In wild-type BALB/c hosts (2-4), relative to Foxn1^nu/nu^ hosts, one would expect depletion of genes required for an effective immune response. Indeed, we observed significant depletion of known mediators of immune evasion (*Cd274/Pd-l1*), components of the IFNγ signaling pathway (*Jak1, Jak2, Stat1* and *Irf1)*, an E3 ubiquitin ligase (*Cop1*), an oncogenic transcriptional activator previously identified by our laboratory in prostate cancer (*Trim24) (Groner et al*., *2016)*, and others (Figure 2C; Table S4). The phenotype of these genes in 4T1 tumors were also observed in a second murine TNBC model (EMT6) (Figures S2G-S2J) and a murine colorectal cancer model (MC38) (Figures S3A-S3D), validating the robustness of our findings.

**Figure 2.**
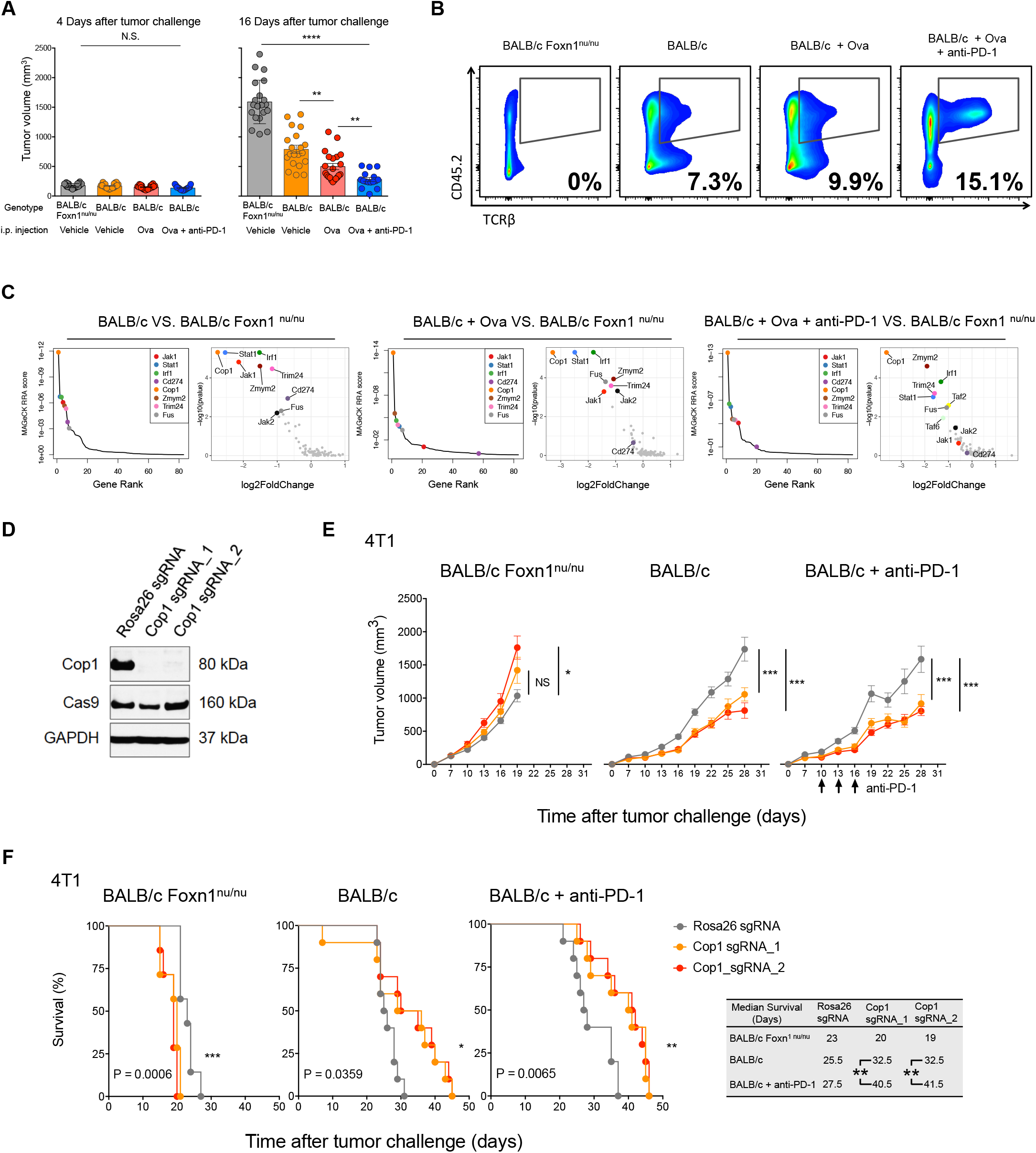
Second Rounds MusCK Screens Identified *Cop1* as a Novel Regulator of TNBC Progression. (A) Tumor volume measured at 4 and 16 days post implantation in the MusCK 2.0 screens. Data are shown as mean ± SEM, n = 7-12 mice per group, **p < 0.01, * * * * P < 0.0001, by one-way ANOVA with Benjamini-Hochberg post-test multiple comparison. (B) Flow cytometry analysis of tumor-infiltrating T cell population (TCRb+) in the total immune cell population (Cd45.2^+^). (C) MAGeCK analysis and RRA ranking of top depleted genes in the MusCK 2.0 screens. Ranked dot plots of depleted genes in immunocompetent hosts compared to immunodeficient nude hosts are shown. (D) Western blot of Cop1 protein level in 4T1 mouse TNBC cells transduced with sgRNA targeting Cop1 and Rosa26. (E) Tumor volume over time in host animals implanted with Rosa26 KO and Cop1 KO 4T1 mouse TNBC cells. Data are shown as mean ± SEM, n = 10 mice per group, *p < 0.05, ***p < 0.001, by one-way ANOVA with Benjamini-Hochberg post-test multiple comparison. (F) Kaplan-Meier survival analysis of host animals bearing Rosa26 and Cop1 KO 4T1 tumors. The sgCop1 cohort with anti-PD-1 treatment survived significantly longer than the other groups. n = 10 mice per group, *p < 0.05, **p < 0.01, ***p < 0.001, by log-rank test.

From our two rounds of *in vivo* screens, *Cop1* emerged as the most significantly depleted gene in 4T1 tumors from immunocompetent mice, relative to nude mice (Figures S3E and S3F). While *Cop1* KO cells did not decrease viability compared to control *Rosa26* KO cells *in vitro* (Figure 2D; Figure S3G), we observed significantly slower tumor progression of *Cop1* KO TNBC cells *in vivo*, in both wild-type BALB/c hosts with and without anti-PD-1 treatment (Figure 2E; Figure S3H). Kaplan-Meier survival analysis showed that mice with *Cop1*-deficient tumors had prolonged survival in wild-type mice, with or without anti-PD-1 treatment, compared to nude mice (Figure 2F). In the MC38 colorectal cancer model of immunocompetent C57BL/6 hosts, *Cop1* KO in cells was able to significantly decrease tumor growth and extend mouse survival (Figures S3I-S3K). Remarkably, in MC38 cells *Cop1* KO together with anti-PD-1 treatment *in vivo* was able to eradicate tumor growth and increase survival to 100% at 60 days post tumor implantation (Figure S3K). The effect of *Cop1* KO in a second, colorectal cancer, syngeneic model suggests that *Cop1* inhibition enhances anti-tumor immunity through a mechanism that may be applicable to other cancer types.

### *Cop1* Increases Macrophage Infiltration by Regulating Macrophage-associated Chemokines

*Cop1* was originally discovered in *Arabidopsis* to induce targeted protein degradation (Osterlund et al., 2000). Multiple substrates of *Cop1-*mediated protein degradation in mammals with cancer implications have been identified, including the classic tumor-suppressor *Tp53* (Dornan et al., 2004a), transcriptional regulator *c-Jun* (Savio et al., 2008; Wertz et al., 2004), and metabolic regulator *Torc2* (Dentin et al., 2007). In humans, *COP1* is located in a region of chromosome 1 frequently amplified in breast cancer patients (Figure S4A) (Dornan et al., 2004b). To characterize the effects of *Cop1* on anti-tumor immunity, we first performed RNA-seq analysis of *Cop1* KO and control *Rosa26* KO in 4T1 cells under IFNγ treatment (Figure 3A). Differential expression analysis showed that 754 genes were significantly up-regulated and 1,303 down-regulated (q < 0.05) upon *Cop1* KO (Figure 3B). Gene set enrichment analysis (GSEA) showed enrichment of downregulated genes in immune-related pathways, including TNFα signaling, inflammatory responses, JAK-STAT signaling pathways, chemokine and cytokine signaling activities (Figure 3C). We also observed similar results in 4T1 cells without IFNγ stimulation (Figures S4B-S4D).

**Figure 3.**
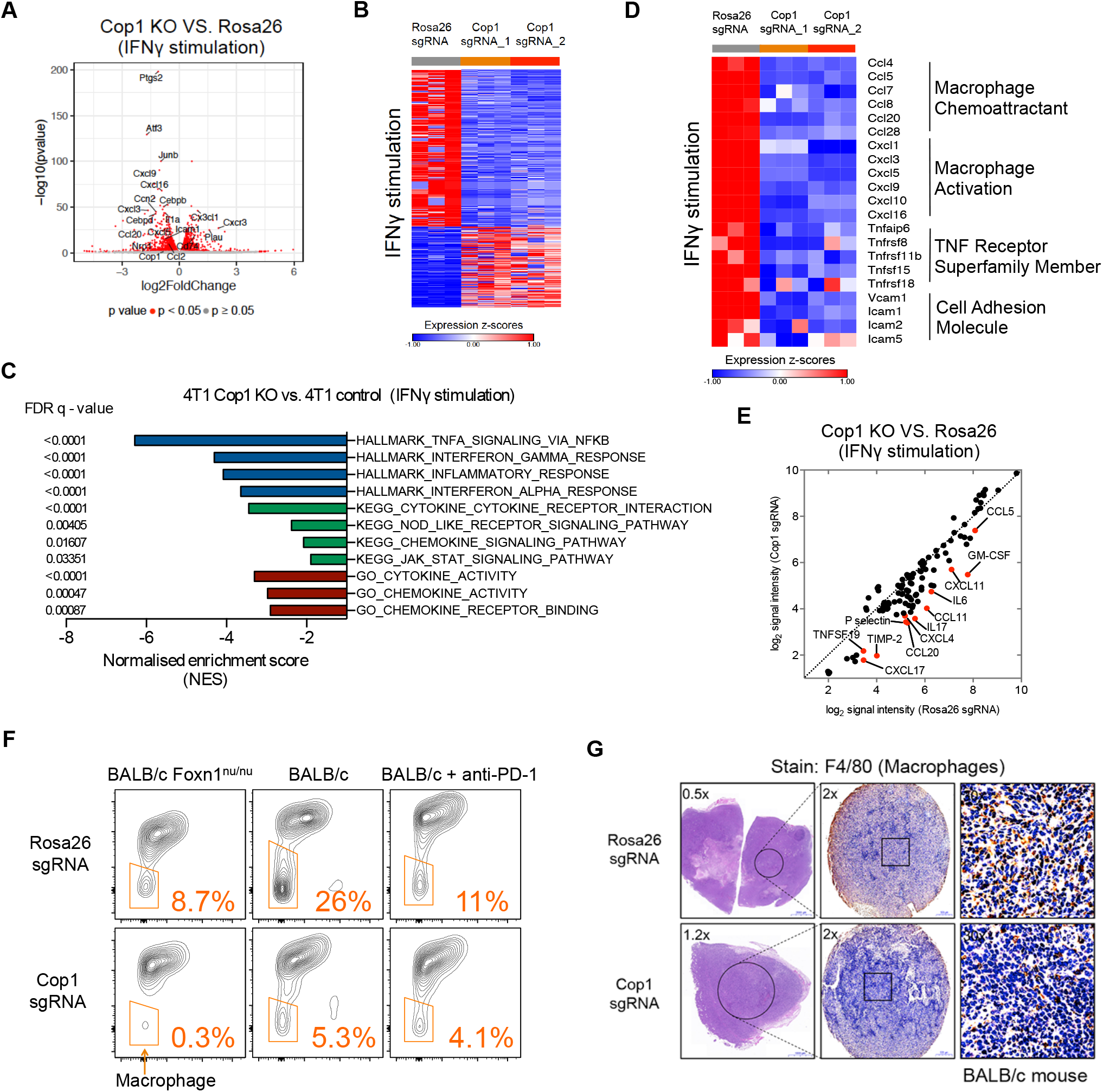
*Cop1* Is a Key Mediator of Macrophage Chemotaxis in TNBC. (A) Volcano plot of differentially expressed genes in Cop1 KO 4T1 mouse TNBC cells compared to Rosa26 KO control cells with IFNγ stimulation (at 20 ng/mL for 24 hours). Red dots denote genes significantly (p < 0.05) differentially expressed in compared conditions. (B) Heatmap showing differential transcriptomic expression in Rosa26 KO and Cop1 KO 4T1 cells with IFNγ stimulation. (C) Gene set enrichment analysis of downregulated genes in Cop1 KO 4T1 cancer cells compared to Rosa26 KO control cells with IFNγ stimulation. Top depleted pathways in Cop1 KO cells versus Rosa26 KO control cells are shown. (D) Differential transcriptomic expression of macrophage-related genes in Rosa26 KO and Cop1 KO 4T1 cells with IFNγ stimulation. (E) Quantification of differential protein expression by cytokine array in Rosa26 KO and Cop1 KO 4T1 cells with IFNγ stimulation. (F) Flow cytometry analysis of macrophage populations in Rosa26 and Cop1 KO 4T1 tumors grown in different host conditions *in vivo*. The tumor-infiltrating macrophages were identified as Cd45.2^+^Cd11c^low^Cd11b^high^Ly6C^low^Ly6G^low^. The tumor-infiltrating myeloid cells were identified as Cd45.2^+^Cd11c^low^Cd11b^high^. (G) Immunohistochemistry of sections show different macrophage infiltration in Rosa26 and Cop1 KO 4T1 tumors. The tumor-infiltrating macrophages were stained by immunohistochemistry with F4/80 antibody, a widely-used monocyte-macrophage marker in mice.

One intriguing result was that *Cop1 KO* in 4T1 cells (Figure S4E), either with or without IFNγ stimulation (Figure 3D), resulted in significant down-regulation of key macrophage chemoattractants, cytokines involved in macrophage activation, and members of the TNF receptor superfamily. Quantification of protein expression based on a cytokine array confirmed significantly decreased levels of cytokines and chemokines known to recruit and activate macrophages, such as *Ccl2, Ccl5, Ccl11, Ccl19, Ccl20, Cxcl4, Cxcl11, Gm-csf*, and *Il-6* (Figure 3E; Figures S4F-S4G). Consistently, flow cytometry and immunohistochemistry found a significant decrease in macrophage infiltration in the *Cop1* KO tumors (Figures 3F-3G; Figures S4H-S4I). In contrast, no significant change was observed in tumor-infiltrating CD8^+^ T cells (Figures S4J and S4K). Furthermore, we confirmed the effect of *Cop1* KO in decreasing macrophage chemoattractants in the tumors grown *in vivo* using cytokine array (Figure S4L). Furthermore, in the 4T1 model, macrophage percentage in tumor-infiltrating Cd45^+^ leukocytes was positively correlated with tumor size, while T-cell percentage was negatively correlated (Figures S4M and S4N). Together, our results suggest that *Cop1* in TNBC regulates macrophage chemotaxis in the TME. Inhibition of *Cop1* decreases tumor macrophage infiltration, which in turn inhibits tumor progression and improves survival.

### Integrated Analyses Identify *C/ebpδ* as a Specific Protein Substrate of *Cop1*

We next sought to identify the putative protein substrates of the E3 ubiquitin ligase *Cop1*. Since most of the known *Cop1* substrates are transcription factors (Dornan et al., 2004a; Migliorini et al., 2011; Vitari et al., 2011), we reasoned that *Cop1* KO might stabilize transcription factor substrates that down-regulate RNA expression of macrophage cytokines. We therefore inferred the likely transcription factors underlying the differentially expressed genes upon Cop1 KO using a computational method that we developed called LISA (Qin et al., 2020). Given a list of differentially expressed genes, LISA uses a large compendium of publicly available ChIP-seq, chromatin accessibility profiles, and transcription factor motifs in the Cistrome database to infer the driving transcription regulators (Mei et al., 2017; Zheng et al., 2019). LISA analysis of the genes downregulated upon *Cop1* KO implicated the CEBP and AP-1 families of transcription factors as putative regulators (Figure 4A). While a function in transcriptional repression has not yet been reported for the CEBP family, the AP-1 family is known to repress gene transcription (Eferl and Wagner, 2003; Miao and Ding, 2003).

**Figure 4.**
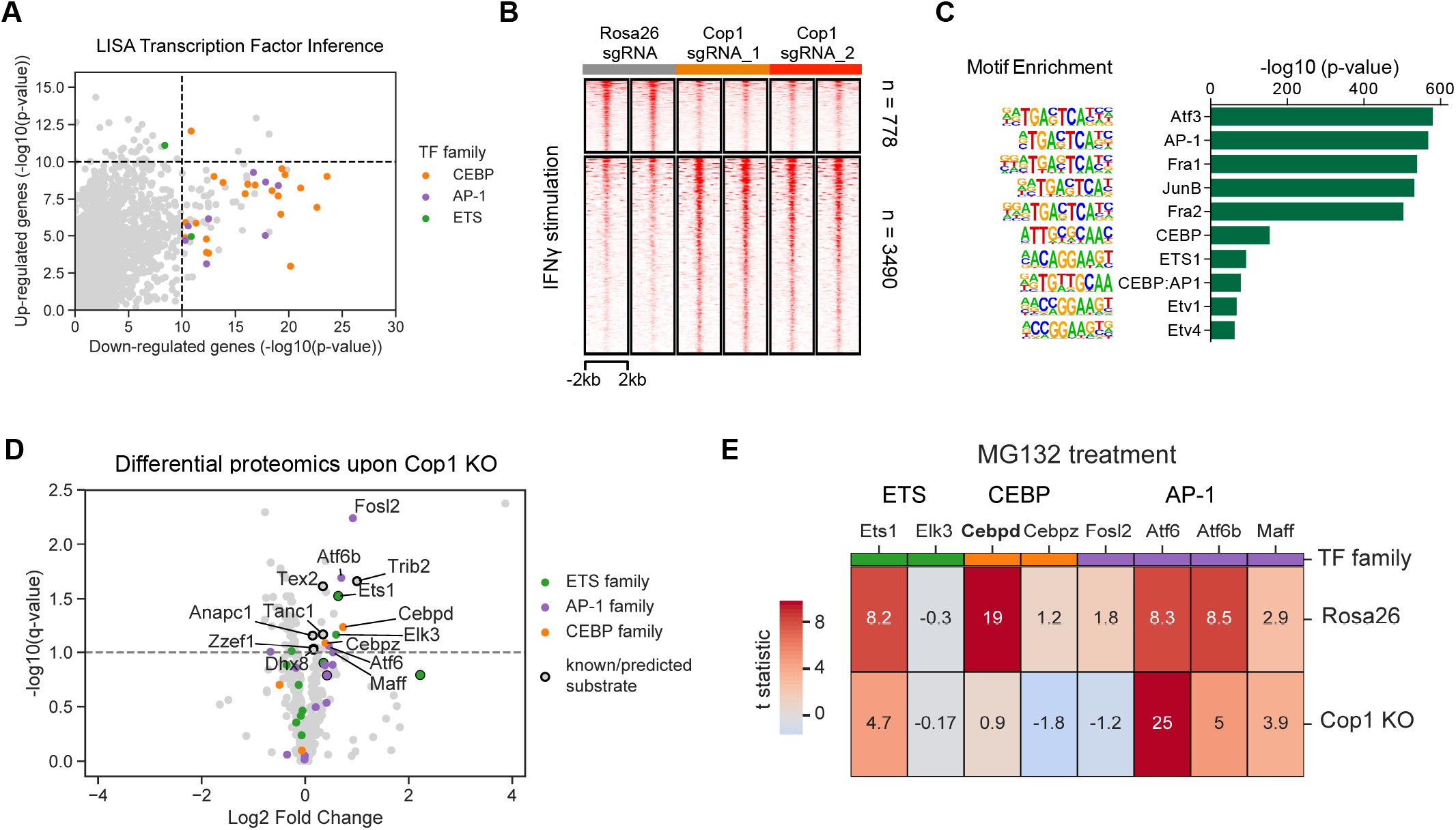
Integrative Analysis Identifies C/ebpδ Activity Is Modulated Upon *Cop1* KO. (A) LISA predicts CEBP and AP1 families of transcription factors in regulating *Cop1* KO down-regulated genes. (B) Heatmap showing changes in chromatin accessibility of Rosa26 and Cop1 KO 4T1 cancer cells with IFNγ stimulation (at 20 ng/mL for 24 hours). (C) Enrichment of known transcription factor motifs in Cop1 KO/Rosa26 differential peaks. (D) Proteomics analysis of Rosa26 and Cop1 KO 4T1 cancer cells. Points above the dashed line are statistically significant (q < 0.1). (E) Heatmap displaying the protein abundance of genes in 4T1 cells with MG132 treatment (proteasome inhibitor). Each row is showing the comparion between proteasome inhibition versus vehicle. If a protein is not degraded by the proteasomal degradation pathway then it should show zero difference in protein expression.

Meanwhile, we hypothesized that the transcription factor substrates stabilized upon *Cop1* KO would in turn increase the chromatin accessibility at the respective transcription factor binding sites. We thus performed ATAC-seq on *Cop1* KO and control Rosa26 KO 4T1 cells. Regardless of IFNγ treatment, *Cop1* KO did not change the overall chromatin accessibility (Figure 4B; Figure S5A), although there were more up-regulated ATAC-seq peaks. An analysis of motif enrichment and peak overlap with public ChIP-seq data found *Cop1* KO-specific up-regulated peaks were enriched for binding by the *Ap-1*, CEBP, and *Ets* families of transcription factors (Figure 4C; Figure S5B; Table S5). Therefore, the ATAC-seq data support RNA-seq analysis in implicating the AP-1 and CEBP families of transcription factors as putative substrates of Cop1 in 4T1 cells.

To further validate the substrates of *Cop1* protein degradation, we used mass spectrometry to find up-regulated proteins in *Cop1 KO* 4T1 cells compared to control cells (Figures S5C and S5D; Table S6). Among the over seven thousand detected proteins, several members of the ETS, AP-1, and CEBP transcription factor families were significantly up-regulated (FDR <0.1), including known *Cop1* substrates *c-Jun, Ets1, Ets2*, and *Etv4* (Figure S5E; Figure 4D). To rule out the possibility of non-proteasomal degradation from secondary effects, we conducted additional proteomics analysis after treating the cells with the proteasome inhibitor MG132. Among the aforementioned families of transcription factors, only *C/ebpδ* showed Cop1-dependent protein degradation with MG132 treatment (Figure 4E). Taken together, these results suggested that in 4T1 cells, *C/ebpδ* is a specific protein substrate of *Cop1*, which likely mediated the increased chromatin accessibility and decreased target gene expressions upon *Cop1* KO.

### *C/ebpδ* Suppresses the Expression of Macrophage Chemokine Genes in Cancer Cells

To gain further insights into the functional impact of *Cop1* KO on *C/ebpδ*, we performed *C/ebpδ* ChIP-seq to map its binding sites and downstream target genes. Consistent with the increased *C/ebpδ* protein abundance upon *Cop1* KO, there was a larger number of up-regulated *C/ebpδ* binding peaks (Figure 5A) and consistently greater chromatin accessibility (Figure 5B), compared to down-regulated peaks. Motif analysis found *CEBP* as the top enriched motif in up-regulated *C/ebpδ* peaks and *AP-1* family member *Fos* as the top motif in down-regulated *C/ebpδ* peaks (Figure 5C), suggesting that the up-regulated *C/ebpδ* peaks are the primary effect of *Cop1* KO on *C/ebpδ*. To assess the effect of the increased *C/ebpδ* binding, we evaluated the differentially expressed genes near the *C/ebpδ* binding sites using the computational tool BETA, which was previously developed in our lab (Wang et al., 2013). BETA found *C/ebpδ* binding sites to be enriched more with down-regulated genes upon *Cop1* KO (Figure S5F), which are significantly associated with regulation of immune-response genes and macrophage chemokines, such as *Ccl2* and *Ccl7* (Figures 5D and 5E). In contrast, *C/ebpδ* binding sites near genes that are up-regulated upon *Cop1* KO are enriched in amino acid metabolism and peptide biosynthesis, suggesting that these represent secondary effects (Figure 5D). Taken together, our results suggest that *Cop1* KO decreased the proteasomal degradation of *C/ebpδ*, and that the stabilization of *C/ebpδ* suppresses transcription of immune response genes and macrophage chemokines.

**Figure 5.**
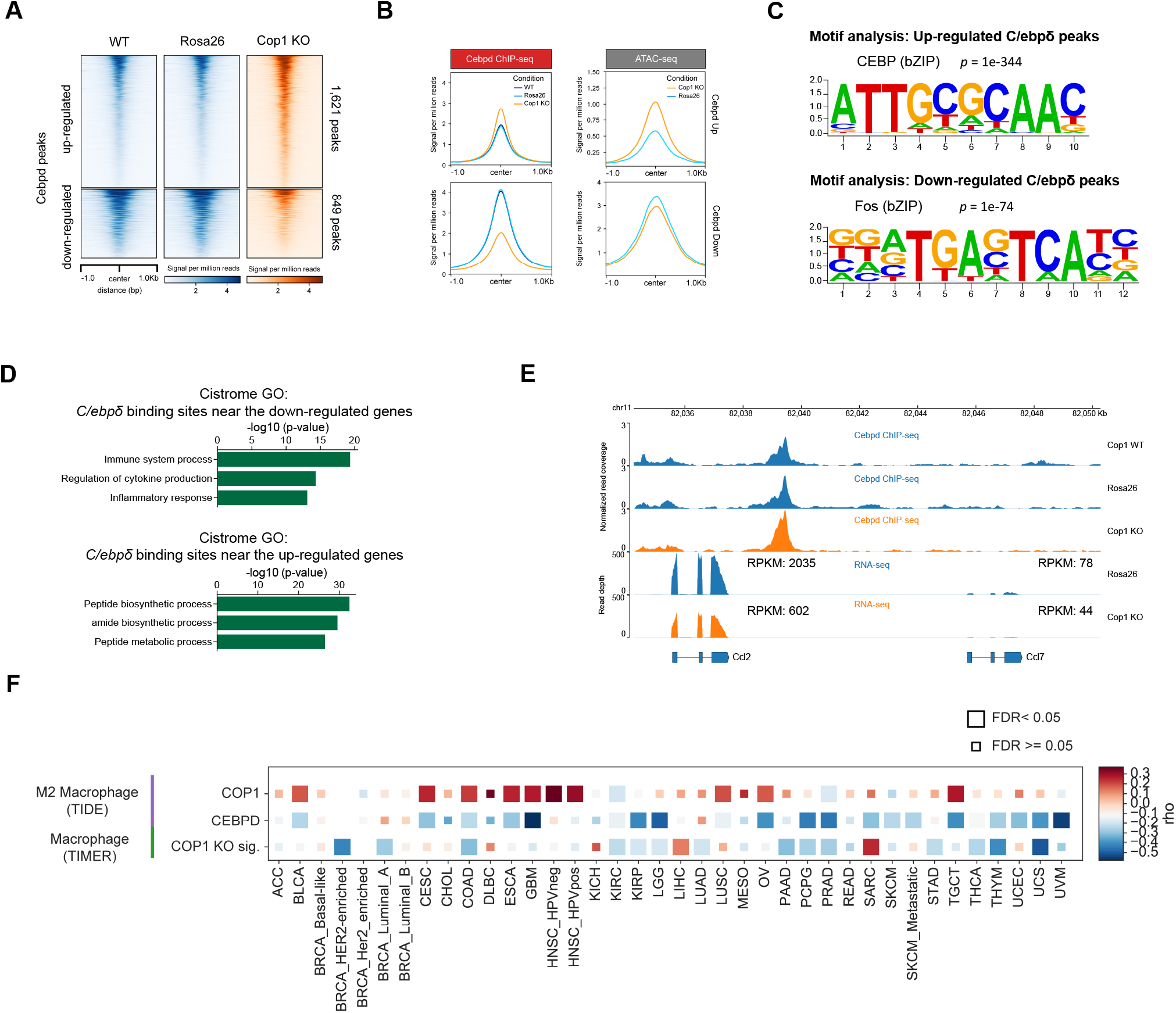
The COP1-axis Is Associated with Macrophage Infiltration and Response to ICB for Cancer Patients. (A) Distribution of normalized read counts in a 2,000 bp window around Cop1 KO-specific C/ebpδ peaks. (B) Distribution of gene-averaged read counts for the datasets of C/ebpδ ChIP-seq and ATAC-seq. (C and D) Significant *de novo* motifs (C) and enriched ontology terms (D) of Cop1 KO-specific C/ebpδ peaks, respectively. p values determined by hypergeometric test. (E) Normalized signal tracks of ChIP-seq and ATAC-seq at the Ccl2 and Ccl7 locus in 4T1 cancer cells. (F) Heatmap showing the correlation between gene expression of COP1 or CEBPD with inferred macrophage infiltration in The Cancer Genome Atlas (TCGA). CEBPD expression was negatively correlated with M2 macrophages (TIDE). Correlations were obtained through the TIMER website and adjusted for tumor purity. Cancer types are labeled on the x-axis.

To evaluate whether the *Cop1* effect on macrophage infiltration and tumor progression in mouse TNBC models (Figures 2E and 3G) is relevant in human tumors, we examined public tumor cohorts. *COP1* is more highly expressed in many cancer types in The Cancer Genome Atlas (TCGA) (Figure S5G), including breast and colon cancers, compared to adjacent normal samples. In addition, lower *COP1* expression in tumors is associated with better outcomes in some cancer types, including breast cancer, ovarian cancer, and papillary kidney cancers (Figures S5H-S5J). Further, *COP1* expression is positively correlated with computationally inferred M2 macrophages, while *CEBPD* expression is negatively correlated with inferred M2 macrophages, across many cancer types (Figure 5F) (Jiang et al). A *Cop1* KO signature of differentially expressed genes (Li et al., 2019, 2017), which may partially reflect *C/ebpδ* protein activity, was also negatively correlated with inferred macrophage infiltration (Figure 5F, see STAR Methods for further details). These results were consistent using different computational algorithms for immune deconvolution analysis (Figures S5K and S5L) (Aran et al., 2017; Newman et al., 2019). Together, these data indicate a robust association of high *Cop1* and low *C/ebpδ* expression with increased macrophage infiltration across human cancers.

### *Cop1* Targets *C/ebpδ* for Proteasome Degradation via the Scaffolding Protein *Trib2*

To elucidate how *Cop1* degrades the *C/ebpδ* protein, we screened proteins that were up-regulated upon *Cop1* KO for a predicted *Cop1* degron motif (Figures S6A-S6C). To this end, we applied a machine learning approach (see STAR Methods for further details) that was predictive of previously reported degrons in known *Cop1* substrates (Figure S6B). This analysis predicted several proteins as the most likely direct *Cop1* substrates in 4T1, including *Trib2 (*Tribbles homolog 2), *Tanc1* (Tetratricopeptide Repeat, Ankyrin Repeat And Coiled-Coil Containing 1), *Tex2* (Testis Expressed 2), and the known substrate *Ets1* (ETS Proto-Oncogene 1) (Figure 4D) (see STAR Methods for further details). Surprisingly, the predicted substrates did not include *C/ebpδ* or any CEBP family member, suggesting that *C/ebpδ* might be an indirect substrate of COP1. We noted that Trib2, the protein with a *Cop1* degron whose level is most elevated upon Cop1 KO, has been previously reported to serve as a substrate adaptor for *Cop1* to modulate its specificity (Figure S6D) (Keeshan et al., 2006). TRIB family pseudokinases possess a C-terminal tail that serves as a peptide motif for MAPKK/MEK family members, and a second binding motif that facilitates direct association with E3 ubiquitin ligases (Eyers et al., 2017). In human or mouse acute myeloid leukemia (AML), TRIB pseudokinases are known to provide a unique molecular scaffold bound by both *C/ebp*α and *Cop1* (Eyers et al., 2017; Jamieson et al., 2018; Murphy et al., 2015). Notably, C/*ebp*δ but not C/*ebp*α was detected at the protein level in 4T1 cells. Based on this, we hypothesized that *Trib2* might serve as an adaptor to facilitate the interaction between *C/ebpδ* and *Cop1*, leading to *Cop1*-mediated proteasomal degradation of *C/ebpδ* (Figure 6A).

**Figure 6.**
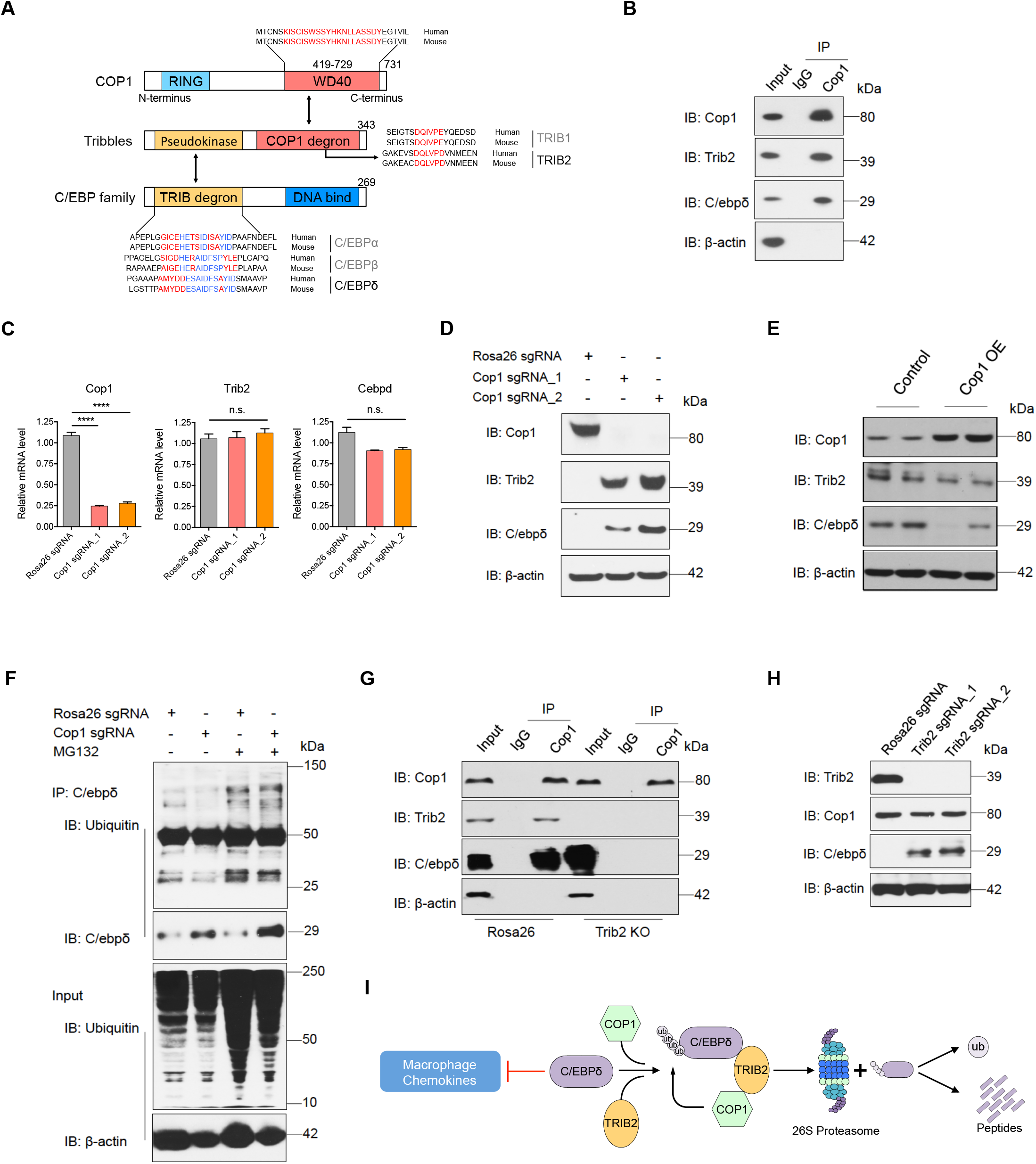
Identification of *C/ebpδ* As a Direct Target of *Cop1* via Adaptor *Trib2*. (A) Schematic illustrating motifs of CEBP family members bound by Tribbles-Cop1. (B) The lysate from wild-type 4T1 cells were incubated with Cop1 antibody or normal control IgG, and the immunocomplexes were probed with the indicated antibodies. (C) Relative mRNA levels of Cop1, Trib2 and C/ebpδ in Rosa26 KO and Cop1 KO cancer cells. Data are shown as mean ± SEM, 3 replicates per condition, ****p < 0.0001, by one-way ANOVA with Benjamini-Hochberg post-test multiple comparison. (D) Western blot showing representative protein levels of Cop1, Trib2 and C/ebpδ in Rosa26 KO and Cop1 KO cancer cells. (E) Western blot showing representative protein levels of Cop1, Trib2 and C/ebpδ in Cop1 overexpressing and control 4T1 cells. (F) Western blot of protein ubiquitination levels of Rosa26 KO and Cop1 KO 4T1 cells under treatment of vehicle or MG132. (G) The lysate from Rosa26 and Trib2 KO 4T1 cells were incubated with Cop1 antibody or normal control IgG, and the immunocomplexes were probed with the indicated antibodies. (H) Western blot showing representative protein levels of Cop1, Trib2 and C/ebpδ in Rosa26 KO and Trib2 KO cancer cells. (I) Schematic illustrating degradation of C/ebpδ by Trib2-Cop1.

To test this hypothesis, we first performed co-immunoprecipitation (Co-IP) in wild-type 4T1 cells to confirm the co-binding of endogenous *Cop1, Trib2*, and *C/ebpδ* (Figure 6B; Figures S6E and S6F). Furthermore, we found that, while *Cop1* KO did not significantly increase *Trib2* and *C/ebpδ* mRNA levels (Figure 6C), it significantly increased their protein levels (Figure 6D). In addition, forced overexpression of *Cop1* led to a decreased *C/ebpδ* protein level without affecting its mRNA (Figure 6E; Figure S6G), supporting *C/ebpδ* post-translational regulation by *Cop1*. To confirm that *C/ebpδ* degradation is mediated by the proteasome pathway, we treated 4T1 cells with selective proteasome inhibitor MG132. We observed that with proteasome inhibition, *Trib2* and *C/ebpδ* proteins were significantly more abundant than in wild-type cells regardless of the *Cop1* status (Figure S6H). Moreover, the polyubiquitination level of *C/ebpδ* was attenuated by *Cop1* KO in the 4T1 cells and elevated with proteasome inhibition (Figure 6F). To prove that *Trib2* is important in mediating *Cop1* degradation of *C/ebpδ*, we used CRISPR to KO *Trib2*. This not only disrupted the interaction between *Cop1* and *C/ebpδ* (Figure 6G), but also increased the protein level of *C/ebpδ* (Figure 6H). Taken together, these results indicate that in 4T1 cancer cells, *C/ebpδ* is a substrate of *Cop1* and the interaction between *Cop1* and *C/ebpδ* is mediated by *Trib2*, which results in ubiquitination and proteasomal degradation of *C/ebpδ* (Figure 6I). *Cop1* inhibition, which stabilizes *C/ebpδ* to suppress macrophage chemoattractant release, can increase tumor sensitivity to immunity and immunotherapy (Figure S6I).

## DISCUSSION

Triple negative breast cancers (TNBC) have immunosuppressive tumor microenvironments (TME), preventing an effective response to checkpoint blockade therapies. There is an urgent need to identify new targets to reprogram the suppressive TNBC TME to enhance immunotherapy efficacies. In this study, we used large-scale *in vivo* CRISPR knockout screens to discover genes that sensitize TNBCs to anti-tumor immunity. We found that the E3 ubiquitin ligase *Cop1* regulates the protein abundance of the transcription factor *C/ebpδ* via an adaptor protein *Trib2. C/ebp*δ transcriptionally suppresses macrophage chemoattractant release from cancer cells. *Cop1* inhibition in TNBCs leads to decreased macrophage infiltration, increased sensitivity to anti-PD-1 treatment, and better survival in mouse models. We also observed association between *COP1* expression, levels of macrophage infiltration, and clinical outcomes in many human cancer types. Our study establishes, for the first time, the role of *Cop1* in modulating macrophage infiltration into cancer cells and suppressing the effects of immunotherapies.

Activation of IFNγ signaling pathway in cancer cells has long been considered to facilitate T-cell antigen recognition and activate T-cell cytotoxicity (Gao et al., 2016). Paradoxically, we found breast cancer cells to be sensitized to immunotherapy by loss-of-function of *Jak1, Stat1*, or *Irf1*, which are downstream effectors of the IFNγ signaling pathway. Supporting our observation, loss of *Jak1* has been reported to prevent progression of breast cancer in mammary cancer models (Chen et al., 2018; Wehde et al., 2018). Studies in breast cancer and melanoma models also found that sustained IFN-γ activation could have the opposite effect from short-term IFN-γ treatment, thus inducing resistance to immunotherapy (Benci et al., 2016; Jacquelot et al., 2019). This may explain why early phase clinical trials of IFN-γ in melanoma patients failed (Meyskens et al., 1990, 1995). Therefore, the anti- and pro-tumor functions of IFNγ might depend on the tumor context, microenvironmental factors, signaling intensity, and signalling duration.

Over the past decade, *Cop1* was found to play an important role in tumor growth and metastasis (Wei and Kaelin, 2011). A number of potential *Cop1* degradation substrates have been identified, including *Tp53, c-Jun, Cebpa, Mek1, p65/RelA, Mkk4, Acc1, Mta1, Foxo1, Torc2*, and *Pea3* (Dornan et al., 2004a; Migliorini et al., 2011; Wei and Kaelin, 2011). With both oncogene and tumor suppressor proteins as putative *Cop1* substrates, characterization of *Cop1* as an oncogene or a tumor suppressor has been inconsistent. Analysis of *COP1* essentiality based on CRISPR screens of hundreds of cancer cell lines in the Dependency Map project shows generally weak effect on cell growth in human cancer cell lines *in vitro* (Tsherniak et al., 2017). This is consistent with the *Cop1* knockout phenotype we observed in mouse breast cancer (4T1) and colorectal cancer (MC38) cells grown *in vitro*. At the same time, *Cop1* KO significantly suppressed tumor growth and prolonged survival in wild-type mice, especially mice treated with immune checkpoint blockade, compared to nude mice. This suggests the effects of *Cop1* on tumor progression to be through TME reprogramming and immune response, thus implicating COP1 as a novel immunotherapy target. However, since *Cop1* is expressed not only in cancer cells but also in immune cells and in normal tissues, we do not rule out the possibility that COP1 might have functions in other cell types or tissues. Future studies, especially if a small molecule inhibitor against *Cop1* is available, are needed to evaluate the systemic effect of *Cop1* inhibition in vivo or in human cancers.

Our study found the biological function of *Cop1* in cancer cells through influencing *C/ebpδ* proteasomal degradation. *C/ebpδ* belongs to the CEBP family of transcription factors, which is known to regulate many biological processes, including cell differentiation, motility, proliferation, cell death, metabolism and immune responses (Ko et al., 2015). Previous study reported that *C/ebp*α stability is critical to prevent *Trib1-Cop1* complex-driven acute myeloid leukemia (Nakamae et al., 2017). Another study found that an aberrant *C/ebp*α protein level resulting from *Trib1* deficiency in haematopoietic cells results in severe reduction of M2-like macrophages in bone marrow, spleen, lung, and adipose tissues (Satoh et al., 2013). A more recent study in Alzheimer’s disease reported that *C/EBP*β in microglia, which drives a potent proinflammatory program, is regulated at the protein level by *COP1 (Ndoja et al*., *2020)*. While these findings are consistent with our observations, we for the first time showed the effect of COP1 on macrophage infiltration and tumor growth through *Trib2* and *C/ebpδ* in solid tumors. In addition, COP1 expression and *Cop1* knockout signature are associated with high and low macrophage infiltration across many human cancer types, respectively. We note that the level of *C/ebpδ* protein is the most significantly changed CEBP family member upon *Cop1* KO, but is not the only CEBP family member whose protein levels are affected. It is possible that in other cancer types or immune cells, *Cop1* KO could stabilize other CEBP family members which function in suppressing macrophage infiltration and tumor growth. In addition, our RNA-seq, ATAC-seq and proteomics analyses suggest that the AP-1 family of transcription factors might interact with *C/ebpδ*, or mediate secondary effects, upon *Cop1* KO. Further studies are needed to pinpoint the specific AP-1 family members involved, and to elucidate this interaction and its effect.

Currently available immune checkpoint blockade antibodies, such as anti-PD-1, anti-PD-L1, and anti-CTLA4, aim to facilitate cancer cell recognition by lymphocytes and increase T cell cytotoxicity. However, the majority of human tumors, especially from breast, prostate, colon, and lung cancers, are tumors with low level of cytotoxic T lymphocytes (CTL) and generally elict low immune activity. Therefore, recent cancer immunology research and immune-oncology drug development have been focused on reprogramming the TME by killing immunosuppressive fibroblasts (Noy and Pollard, 2014) or macrophages (Motz and Coukos, 2013) to help T-cell infiltration. The fact that different syngeneic tumor models have very different TMEs indicates that cancer-cell intrinsic mechanisms may determine whether a TME supports an effective or ineffective immune. Our study, together with previous work (Lawson et al., 2020; Manguso et al., 2017), demonstrates the effectiveness of in vivo CRISPR screens in identifying such cancer-cell intrinsic TME regulators. These in vivo studies could only test a limited number of genes in a limited number of syngeneic tumor models. We foresee similar approaches being applied to more genes in more syngeneic models to identify additional targets that can reprogram the TME to enhance immunotherapy response.

## ACKNOWLEDGMENTS

We acknowledge Jin Zhao and Jingyu Peng for their assistance and discussions. This study was supported by the Breast Cancer Research Foundation BCRF-20-100 (to X.S.L.), NIH Grants R01CA234018 (to X.S.L.), Breast Cancer Research Foundation (to M.B.), the Damon Runyon Cancer Research Foundation DRQ-04-20 (to C.T.), and the Sara Elizabeth O’Brien Trust (to S.S.G.).

## AUTHOR CONTRIBUTIONS

X.S.L., M.B. and X.W. conceived, designed, and initiated the study. X.W., S.S.G., N.T. and T.X. generated lentiCRISPR vectors and libraries (MusCK 1.0 and 2.0). X.W., S.S.G. and N.T. performed most *in vivo* experiments. X.W. and N.T. performed cell competition assays. C.T., B.W., W.L., and P.J. analyzed CRISPR screen data. B.W. analyzed mRNA-seq and ATAC-seq data. C.T. analyzed ChIP-seq and proteomics data. X.W. and Y.L. performed biochemistry experiments. B.Z., Z.L. and K.L. assisted X.W. for various experiments. N.T. assisted X.W. for histology experiments. P.C. and H.L. provided suggestions for ATAC-seq experiment and data analysis. C.M., Y.Z. and J.F. provided suggestions for ChIP-seq and single cell RNA-seq data analysis. X.S.L., M.B., X.W., C.T. and S.S.G. jointly prepared the manuscript with inputs from all authors. X.S.L. and M.B. secured funding and supervised the work.

## DECLARATION OF INTERESTS

M.B. is a consultant to and receives sponsored research support from Novartis. MB is a consultant to and serves on the scientific advisory boards of Kronos Bio, H3 Biomedicine, and GV20 Oncotherapy.

X.S.L. is a cofounder, board member, SAB member, and consultant of GV20 Oncotherapy and its subsidiaries; SAB of 3DMed Care; consultant for Genentech; and stock holder of BMY, TMO, WBA, ABT, ABBV, and JNJ, and received research funding from Takeda and Sanofi.

## Supplementary Figure Legends

**Figure S1. Related to Figure 1. Optimization and Validation of Primary MusCK *In Vivo* CRISPR Screens in the Mouse 4T1 TNBC Models**.

(A) Pie chart showing the fraction of genes targeted in the MusCK library.

(B) Protein Cas9 expression in MusCK library-infected 4T1 cells.

(C) Expression of hormone receptors and PD-L1 in mouse and human cancer cells.

(D) Principal component analysis of transcriptome of mouse and human breast cancer cells.

(E) MAGeCK analysis and RRA ranking of top depleted genes in the in vitro MusCK screen.

(F) MAGeCK analysis and RRA ranking of top enriched genes in the in vitro MusCK screen.

(G) Mouse 4T1 cell line was transduced with a vector expressing various forms of ovalbumin antigen.

(H) Western blot of 4T1 cell lysate for ovalbumin and GAPDH after transfection with either control or ovalbumin vector.

(I) Tumor volume of wild-type or ovalbumin expressing 4T1 cancers.

(J) Tumor infiltrating T cells in wild-type or ovalbumin expressing 4T1 cancers.

(K) Western blot of 4T1 cell lysate for Cas9, ovalbumin and GAPDH after transduction with CRISPR library.

(L) Flow cytometry analysis of T cells and B cells in different host conditions.

(M) Tumor volume averaged for groups indicated and tumor weight measured at 16 days post implantation. n = 12 per group.

(N) Gene set enrichment analysis of 4T1 tumor and 4T1 cells. Top enriched pathways in 4T1 tumors versus 4T1 cells were shown. GSEA terms significantly upregulated in 4T1 tumors compared with 4T1 cells.

(O) Scheme of *in vivo* competition between *Jak1* or *Stat1* KO and control *Rosa26* KO cells.

(P) Western blotting of *Jak1* and *Stat1* expression in 4T1 and EMT6 cells transduced with single gRNA targeting *Jak1 (*or *Stat1)*.

**Figure S2. Related to Figure 2. MusCK 2**.**0 Library Screens on 4T1 and EMT6 TNBC Mouse Models**.

(A) Scheme of MusCK 2.0 *in vivo* screen.

(B) Tumor volume of 4T1 tumor averaged for groups indicated and tumor weight measured at 16 days post implantation. Error bars represent +/-1 SEM, n = 10 mice per group.

(C) A matrix of the Pearson’s correlations of the library distribution from one animal compared to any other animal for MusCK 2.0 screen using 4T1 cells.

(D) Representative plots showing the gating strategy for different populations of tumor infiltrating immune cells.

(E and F) Flow cytometry analysis of tumor infiltrating immune cell population in mice under different treatment. Data are shown as mean ± SEM, n = 5 per group, *p < 0.05, **p < 0.01, ***p < 0.001, ****p < 0.0001, by one-way ANOVA with Benjamini-Hochberg post-test multiple comparison.

(G) Tumor volume of EMT6 tumor averaged for groups indicated. Error bars represent +/-1 SEM, n = 10-12 mice per group.

(H) EMT6 tumor volume and weight measured at 16 days post implantation. Data are shown as mean ± SEM, n = 10-12 mice per group, *p < 0.05, ****p < 0.0001, by one-way ANOVA with Benjamini-Hochberg post-test multiple comparison.

(I) A matrix of the Pearson’s correlations of the library distribution from one animal compared to any other animal for MusCK 2.0 screen using EMT6 cells.

(J) MAGeCK analysis and RRA ranking of top depleted genes in the MusCK 2.0 screen.

**Figure S3. Related to Figure 2. MusCK 2**.**0 Library Screens on MC38 Colon Cancer Mouse Models and Identification of Cop1 Function on Mouse Models**.

(A) Tumor volume of MC38 tumor averaged for groups indicated. Error bars represent +/-1 SEM, n = 10-12 mice per group.

(B) Tumor volume and weight measured at 19 days post implantation. Data are shown as mean ± SEM, n = 10-12 mice per group, *p < 0.05, ****p < 0.0001, by one-way ANOVA with Benjamini-Hochberg post-test multiple comparison.

(C) A matrix of the Pearson’s correlations of the library distribution from one animal compared to any other animal for MusCK 2.0 screen using MC38 cells.

(D) MAGeCK analysis and RRA ranking of top depleted genes in the MusCK 2.0 screen.

(E) Frequency of sgRNAs targeting Cop1 in the MusCK 2.0 screen on 4T1 and EMT6 TNBC mouse models.

(F) Behavior of individual sgRNAs targeting Cop1 in MusCK 2.0 screen in 4T1 TNBC models.

(G) Cell viability of *Rosa26* and *Cop1* KO 4T1 cells cultured *in vitro*. Data are shown as mean ± SEM, n = 6 per group, by one-way ANOVA with Benjamini-Hochberg post-test multiple comparison.

(H) Flow cytometry analysis of *Cop1* KO cells versus control cells in the resulting 4T1 tumors.

(I) Flow cytometry analysis of *Cop1* KO cells versus control cells in the resulting MC38 tumors.

(J and K) Tumor volume and Kaplan-Meier survival analysis of host animals bearing Rosa26 and Cop1 KO MC38 tumors under immunoglobulin G (IgG) isotype or anti-PD-1 antibody treatment. Data are shown as mean ± SEM, n = 10 mice per group, *p < 0.05, ***p < 0.001, by one-way ANOVA with Benjamini-Hochberg post-test multiple comparison.

**Figure S4. Related to Figure 3. *Cop1* Knockout Sensitized Cancer Cells to Immune-mediated Cytotoxicity**.

(A) Location of *COP1* (*RFWD2*) gene on chromosome 1 and percentage of subjects with *COP1* gene amplification (red) in different breast cancer data sets.

(B) RNA-sequencing of *Rosa26* and *Cop1* KO 4T1 cancer cells without IFNγ stimulation. Red dots denote genes significantly (p < 0.05) differentially expressed in conditions compared.

(C) Heatmap showing differential transcriptomic expression in *Rosa26* and *Cop1* KO 4T1 cells without IFNγ stimulation.

(D) Gene set enrichment analysis of *Rosa26* and *Cop1* KO 4T1 cancer cells without IFNγ stimulation. Top depleted pathways in *Cop1* KO cells versus *Rosa26* control cells were shown.

(E) Differential transcriptomic expression of macrophage-related genes in *Rosa26* and *Cop1* KO 4T1 cells without IFNγ stimulation.

(F) Representative images of chemokines from proteome array analysis of control (*Rosa26*) and *Cop1* KO 4T1 in vitro cell culture supernatants without IFNγ stimulation.

(G) Representative images of chemokines from proteome array analysis of control (*Rosa26*) and *Cop1* KO 4T1 in vitro cell culture supernatants with IFNγ stimulation.

(H) Quantification of percentage of macrophages in all tumor-infiltrating myeloid cells. Data are shown as mean ± SEM, n = 8-10 per group, *p < 0.05, **p < 0.01, ***p < 0.001, by one-way ANOVA with Benjamini-Hochberg post-test multiple comparison.

(I) Immunohistochemistry of sections showing monocyte chemoattractant CCL2/MCP1 expression in *Rosa26* and *Cop1* KO 4T1 tumors.

(J) Flow cytometry analysis of CD8^+^ T cell populations in *Rosa26* and *Cop1* KO 4T1 tumors grown in BALB/c mice under different treatment *in vivo*. Quantification of percentage of CD8^+^ T cells in all tumor-infiltrating T cells.

(K) Immunofluorescence of sections showing different T cell infiltration in *Rosa26* and *Cop1* KO 4T1 tumors.

(L) Representative images of chemokines from proteome array analysis of control (*Rosa26*) and *Cop1* KO 4T1 tumors. Quantification of differential protein expression (in cytokine array) of tumor extracts harvested from host animals.

(M) Correlation of macrophage infiltration with tumor volume at end point.

(N) Correlation of T cell infiltration with tumor volume at end point.

**Figure S5. Related to Figures 4 and 5. Prioritization of Putative Cop1 Substrates**.

(A) Heat map showing changes in chromatin accessibility of Rosa26 and Cop1 KO 4T1 cancer cells without IFNγ stimulation.

(B) TF motif enrichment of Rosa26 and Cop1 KO 4T1 cancer cells with or without IFNγ stimulation. Expected (x axis) versus observed (y axis) percentages of IFNγ treated cells peaks overlapping each TF binding site annotation.

(C) Western blot of protein ubiquitination levels in Rosa26 KO and Cop1 KO 4T1 cells under treatment of vehicle or MG132 (at 50 μM for 8 hours).

(D) Heatmap displaying the correlation in the proteome for samples indicates consistency among replicates. Color indicates the spearman correlation.

(E) Differential protein abundance (Cop1 KO VS. Rosa26 4T1 cells) of previously reported substrates of Cop1.

(F) BETA analysis of C/ebpδ binding sites in Cop1 KO and Rosa26 KO 4T1 cancer cells.

(G) Compared to adjacent normal tissue, COP1 was overexpressed in TCGA cancer samples. Data are shown as mean ± SEM, *p < 0.05, ***p < 0.001, by Mann-Whitney U test.

(H) Kaplan-Meier plot displaying that higher COP1 expression associates with worse survival in the ovarian and kidney cancer cohorts (Cox PH test).

(I and J) Kaplan-Meier plot displaying that higher COP1 expression associates with worse survival in the METABRIC breast cancer cohort (Cox PH test). The association with survival was still significant when restricting Triple Negative Breast Cancer (TNBC) (Mann-Whitney U test).

(K-L) Partial spearman correlation after adjusting for tumor purity of the Cop1 gene expression signature with estimated immune cell infiltration of tumors from J) Cibersort, and K) xCell. X-axis reflects cancer types from TCGA and y-axis is immune cell types.

**Figure S6. Related to Figure 6. Trib2 Serve as a Substrate Adaptor for Cop1 to Target C/ebp**δ.

(A) Diagram displaying that putative Cop1 substrates must be up-regulated at the protein-level upon Cop1 KO, contain a Cop1 degron motif, and score well according to a machine learning model.

(B) Scoring of the degron-likelihood by the RF algorithm of Cop1 motifs indicates known Cop1 substrates received higher scores. Box plot represents quartiles +/-1.5 interquartile range. Mann-Whitney U test.

(C) Overlapping the results from mouse and human showed 117 motif hits as possible Cop1 degrons (score>0.5).

(D) Protein interaction network of human COP1 (Bioplex database) indicates tribbles protein family (TRIB1/2/3) physically interact with COP1. Colors denote differential expression upon Cop1 KO: red=up-regulated, blue=down-regulated.

(E and F) The lysate from wild-type 4T1 cells were incubated with Trib2 and C/ebpδ antibody or normal control IgG, and the immunocomplexes were probed with the indicated antibodies.

(G) Real time PCR analysis confirmed that the Cop1 overexpression condition was indeed overexpressed compared to the empty vector.

(H) Western blotting showing representative protein levels of Cop1, Trib2 and C/ebpδ in Cop1 KO and Rosa26 KO 4T1 cancer cells with or without MG132 treatment.

(I) Model for Cop1-driven macrophage infiltration.

## Supplemental Table Legends

Table S1: Guide RNA design for MusCK library.

Table S2: *In vitro* MusCK library screening on 4T1 TNBC cells.

Table S3: *In vivo* MusCK library screening on 4T1 TNBCcells.

Table S4: *In vivo* MusCK2.0 library screening on 4T1 TNBC cells.

Table S5: ATAC-seq analysis of *Cop*1 KO and Rosa26 KO 4T1 cells.

Table S6: Proteomics of *Cop1* KO and Rosa26 KO 4T1 cells.

Table S7: Primers for library construction, real-time PCR and CRISPR KO.

## STAR+METHODS

### KEY RESOURCES TABLE

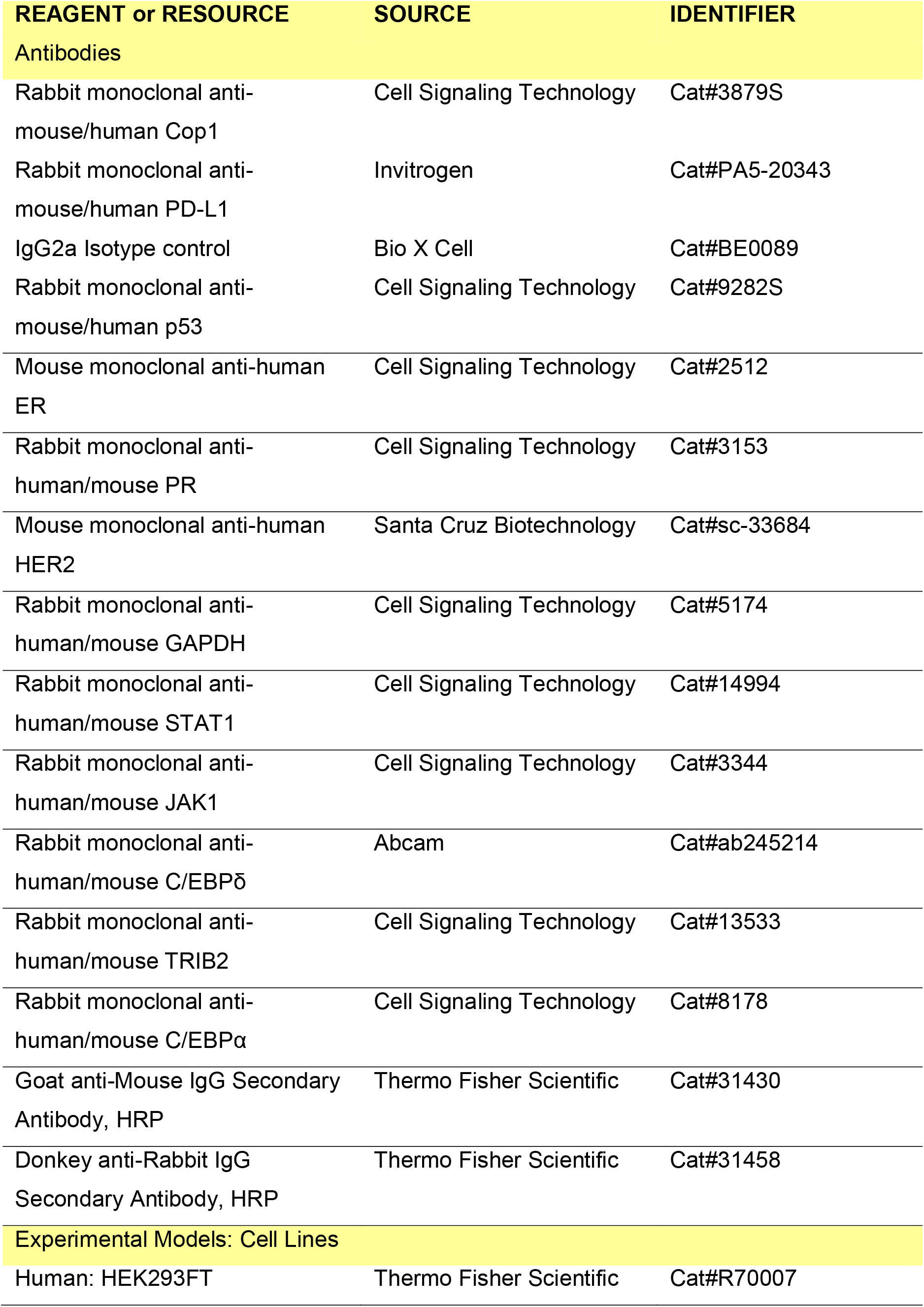
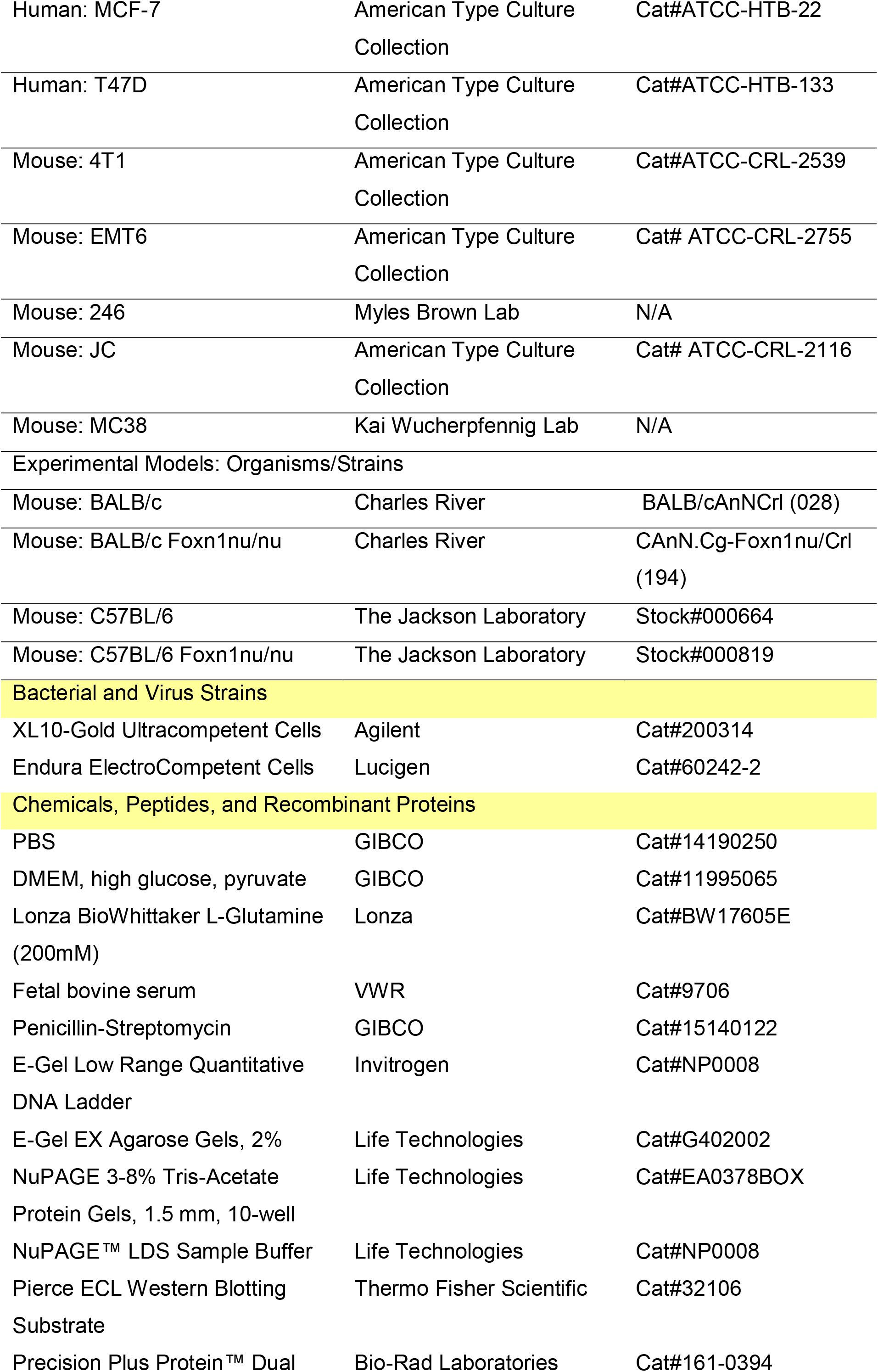
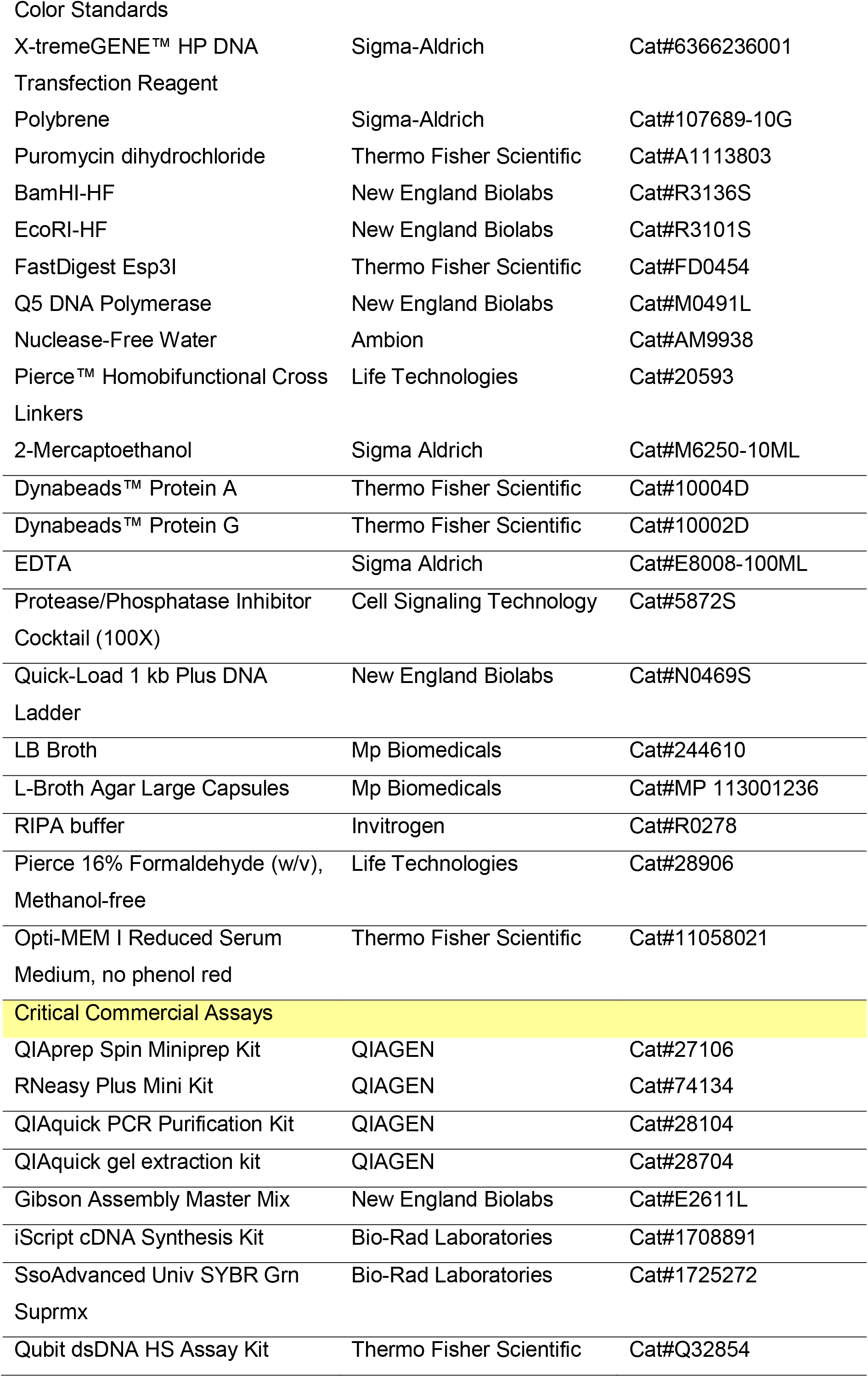
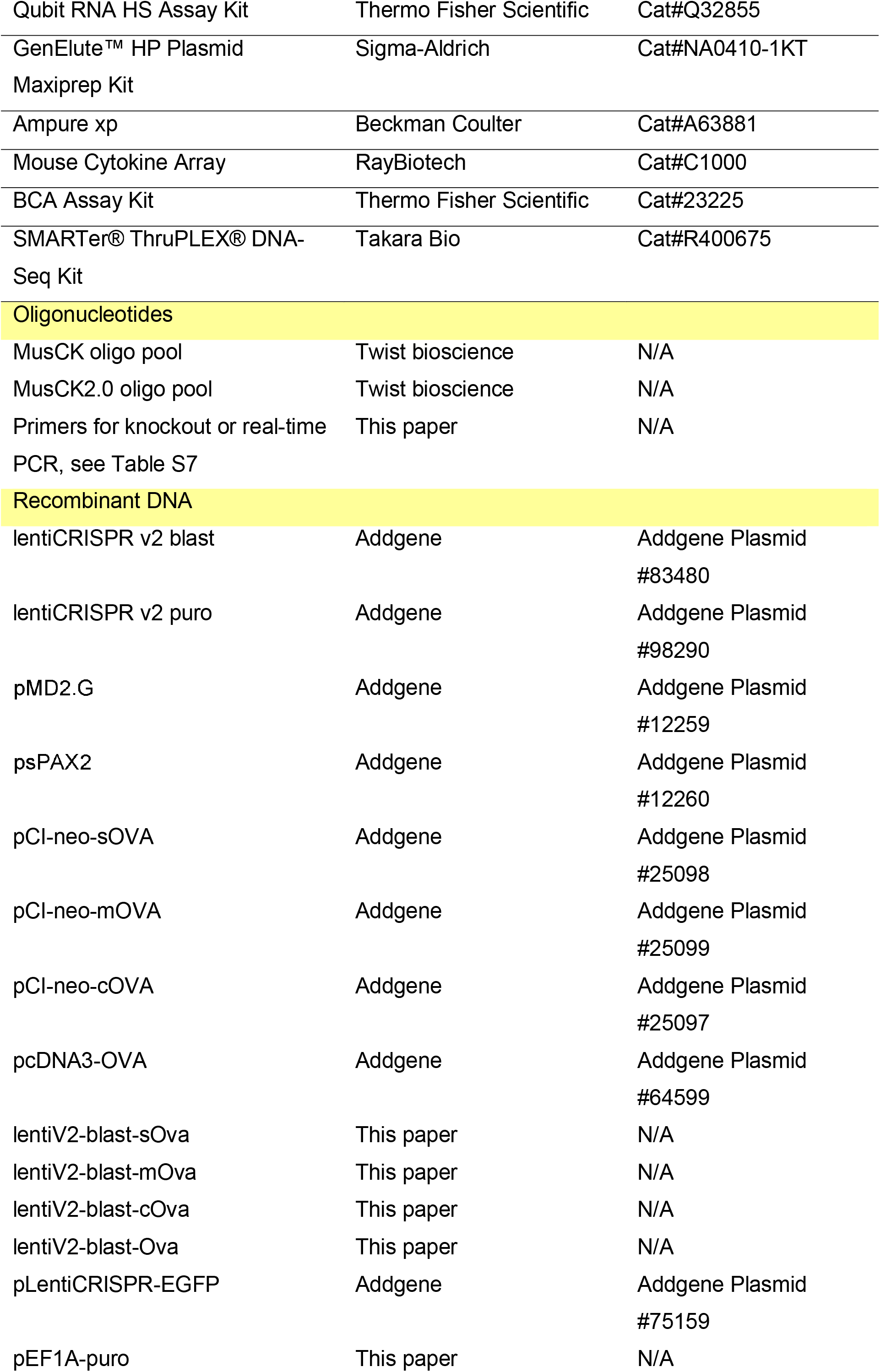
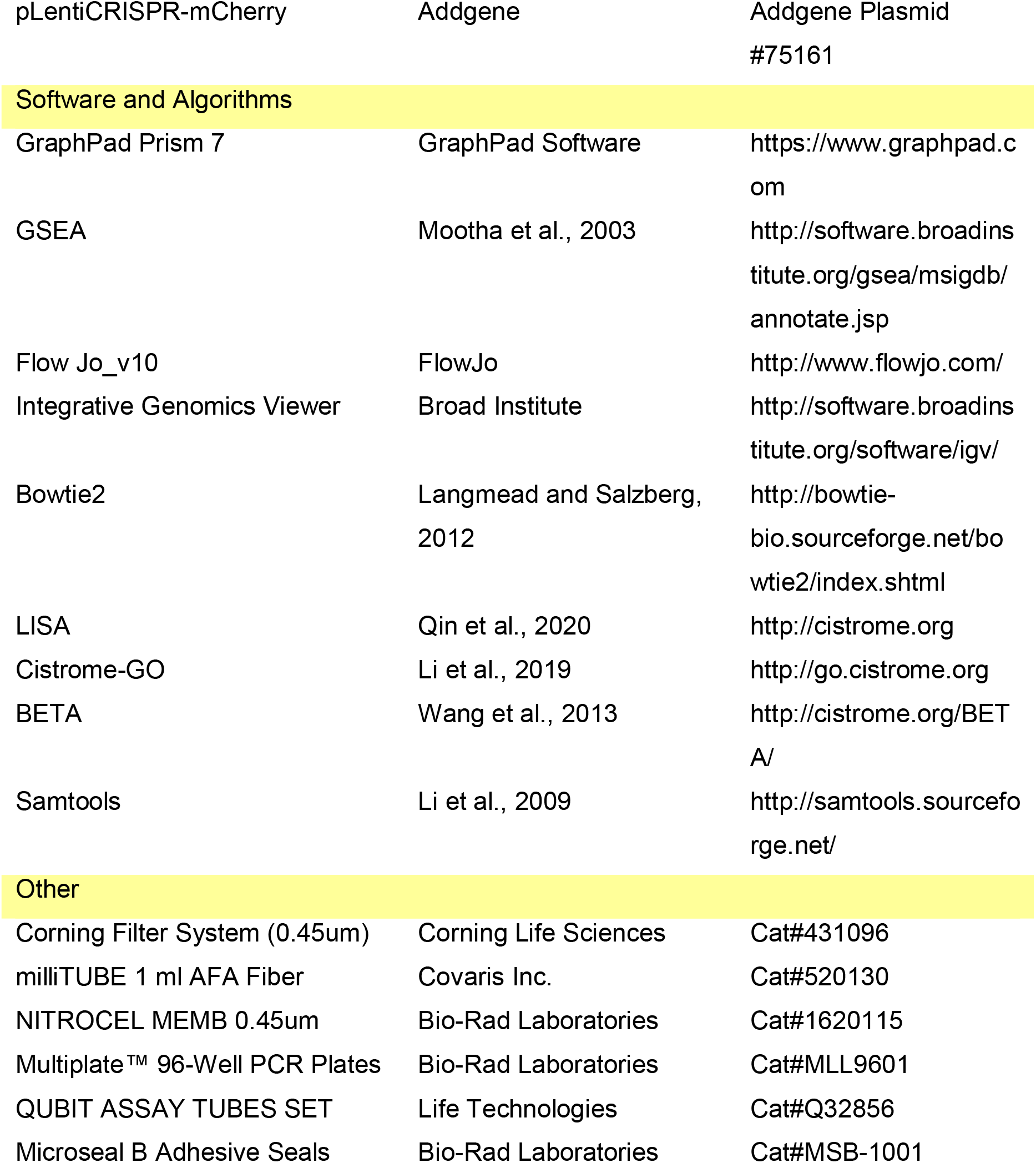

### CONTACT FOR REAGENT AND RESOURCE SHARING

Further information and requests for reagents may be directed to, and will be fulfilled by, the Lead Contact Myles Brown. A list of critical reagents (key resources) is included in the Key Resources Table. Relevant plasmids are available to the academic community. For additional materials, please email the lead contact for requests. Some material may require requests to collaborators and/or agreements with various entities. Requests are reviewed by DFCI regarding intellectual property or confidentiality obligations. Material that can be shared will be released via a Material Transfer Agreement.

### EXPERIMENTAL MODEL AND SUBJECT DETAILS

#### Mice

All animal experiments were approved by the Institutional Animal Care and Use Committee (IACUC) of Dana Farber Cancer Institute (DFCI) and performed with approved protocol (16-005). Six to eight week old female BALB/c (Stock# 028) and BALB/c Foxn1^nu/nu^ (Stock# 194) were obtained from Charles River Laboratory (Wilmington, MA). Six to eight week old female C57BL/6 (Stock# 000664) and C57BL/6 Foxn1^nu/nu^ (Stock# 000819) were obtained from Jackson Laboratory (Bar Harbor, ME). All animals were housed in standard individually ventilated, pathogen-free conditions, with 12h : 12h light cycle, room temperature (21-23°C) and 40%-60% relative humidity. When a cohort of animals were receiving multiple treatments, animals were randomized by 1) randomly assign animals to different groups using littermates, 2) random mixing of mice prior to treatment, maximizing the evenness or representation of mice from different cages in each group, and/or 3) random assignment of mice to each group, in order to minimize the effect of gender, litter, small difference in age, cage, housing position, where applicable. Average tumor sizes were consistent between treatment groups to account for selection bias.

#### Cell Lines

Murine 4T1, EMT6, JC breast cancer cells were obtained from American Type Culture Collection (ATCC) and cultured according to standard protocols. MCF7 and T47D human breast cancer cells were derived and cultured as previously described (Xiao et al., 2018). MC38 murine colon adenocarcinoma cells were obtained from Kai Wucherpfennig laboratory.

## METHOD DETAILS

### Large-scale mouse CRISPR library cloning

SgRNA design primarily targeted low G-C content regions of the genome. We assigned predicted performance scores to all possible sgRNAs targeting each gene, and selected top candidate sgRNAs with the highest predicted on-target KO efficiency and lowest off-target efficiency (Chen et al., 2018a; Xu et al., 2015). Customized single-stranded oligonucleotide pools of CRISPR guide RNA (sgRNA) libraries were synthesized by Twist Bioscience (South San Francisco, CA). The double-stranded oligonucleotides were generated by polymerase chain reaction and cloned into lentiviral CRISPR vector (lentiCRISPR-v2-puro) by Gibson assembly at estimated equal molar ratios to generate the large-scale mouse CRISPR library (MusCK and MusCK 2.0 libraries). The MusCK library consisted of 24,622 sgRNAs including 1,000 non-targeting controls (NTCs) and 23,622 unique sgRNAs targeting 4,787 gene locations in the genome. The MusCK 2.0 library consisted of 800 sgRNAs including 168 non-targeting controls (NTCs) and 632 unique sgRNAs targeting 79 gene locations in the genome. We were aware that in large-scale CRISPR screen efforts, the statistical power of discovery is particularly sensitive to the behavioral consistency of multiple sgRNAs for each target gene. “Outlier” behavior (extreme depletion or enrichment) of one sgRNA out of all sgRNAs targeting the same gene could result in a false positive result. We recognize that the CRISPR KO libraries designed by the Broad Institute is so far the most widely accepted in genomic screen studies; thus, we wanted to ensure that our findings by the MusCK library is reproducible when the Broad sgRNA design principles were applied. To this end, in the MusCK 2.0 library, eight sgRNAs (four designed by our group in MusCK, another four referenced from the Broad Institute’s Brie Mouse CRISPR Knockout Pool Library (Doench et al., 2016) were designated to each candidate gene. An estimated library coverage of ∼300X (total colonies / sgRNAs) was achieved by electroporation. These libraries were subsequently sequence-verified by Illumina sequencing to ensure the high quality of sgRNA distribution.

### Viral library production

The CRISPR library plasmids were transfected into HEK293FT cells at 90% confluence in 15cm tissue culture plates. Viral supernatant was collected at 48 hours and 72 hours post-transfection, filtered via a 0.45 μm filtration unit (Corning, Cat# 430770). The supernatant was subsequently aliquoted and stored in -80 °C freezer until use.

### Viral transduction of cancer cells

Cancer cells were cultured according to standard protocols. Similar to our previous studies (Fei et al., 2017; Xiao et al., 2018), for the pooled large-scale CRISPR screen, a total of > 1×10^8^ cancer cells were transduced with lentivirus containing the library described above at a multiplicity of infection (MOI) of ∼0.3. After puromycin selection for 3 days, ∼30% of the surviving cells were stored as Day-0-input samples at -80°C, and the rest of cells were cultured for in vitro or in vivo screenings. PCR of the regions targeted by the library was performed on genomic DNA to construct the sequencing library. Each library was sequenced at ∼30 million reads to achieve ∼300-fold coverage over the CRISPR library. Sequencing data were analyzed by using MAGeCK and MAGeCK-VISPR (Li et al., 2014, 2015; Wang et al., 2019).

### Genomic DNA extraction

For genomic DNA extraction, two methods were used. Method 1: for cellular samples with a total number greater than 3 ×10^7^ cells, or tumor samples from mice, a custom DNA extraction protocol was used. Briefly, frozen tumors were disrupted on dry ice, then resuspended in 7 mL of Lysis Buffer (400 mM Sodium chloride 10 mM Tris, 2 mM EDTA, 0.5% SDS, pH 8) in a 15 ml conical tube, and 80 μL of 20 mg/ml Proteinase K (Invitrogen) were added to the tumor/cell samples and incubated at 55°C for at least 6 hours. The next day, 80 μl of 20 mg/ml RNase A (Invitrogen) was added to the lysed sample, which was then inverted 10 times and incubated at 65°C for 60 minutes. Samples were cooled on ice before addition of 7 mL of pre-chilled phenol/chloroform (Ambion) to precipitate proteins. The samples were vortexed at high speed for 20 seconds and then centrifuged at 14,000 rpm for 10 minutes. Then, the upper aqueous phase was carefully decanted into a new 15 mL conical tube. Then 7 ml freshly prepared 70% ethanol was added to the tube, vortexed at high speed for 20 second and centrifuged at 12,000 rpm for 10 minutes. Genomic DNA was visible as a small white pellet in each tube. The supernatant was discarded, 6 ml of 70% ethanol was added, the tube was inverted 10 times, and then centrifuged at 12,000 rpm for 5 minutes. The supernatant was discarded by pouring; the tube was briefly spun, and remaining ethanol was removed using a P200 pipette. After air-drying for more than 30 minutes, the DNA changed appearance from a milky white pellet to slightly translucent. Then, 500 μl of nuclease-free water was added, the tube was incubated at 4°C overnight to fully resuspend the DNA. The next day, the gDNA samples were vortexed briefly. The gDNA concentration was measured using a Nanodrop (Thermo Scientific). Method 2: for cellular samples with a total number < 1 ×10^7^ cells, samples were subjected to Allprep DNA/RNA Mini Kit (QIAGEN) following the manufacturer’s protocol.

### sgRNA library readout by deep sequencing

The sgRNA library readout was performed using a two-steps PCR strategy, where the first PCR includes enough genomic DNA to preserve full library complexity and the second PCR adds appropriate sequencing adapters to the products from the first PCR.

For PCR#1, a region containing sgRNA cassette was amplified using primers specific to the lentiCRISPR-v2 vector (Primers for sequencing library construction, see Table S7). PCR was performed using Q5 High-Fidelity DNA Polymerase (NEB). For reactions using Q5 High-Fidelity DNA Polymerase, in PCR#1, the thermocycling parameters were:

| STEP | TEMP | TIME |
| --- | --- | --- |
| Initial Denaturation | 98°C | 30 seconds |
| 18 Cycles | 98°C | 15 seconds |
|  | 68°C | 25 seconds |
|  | 72°C | 25 seconds |
| Final Extension | 72°C | 2 minutes |
| Hold | 4°C |  |

In each PCR#1, we used a different amount of gDNA per sample to capture the full representation of the screen. For example, ∼300X coverage of our genome-wide sgRNA library, gDNA from 3 x 10^7^ cells was used, assuming 6.6 µg of gDNA for 10^6^ cells, 200 µg of gDNA was used per sample (6-8 µg of gDNA per reaction). PCR#1 products for each biological sample were pooled and used for amplification with barcoded second PCR primers (Supplementary Tables). For each sample, we performed at least 3 PCR#2 reactions using 2 µL of the pooled PCR#1 product per PCR#2 reactions for 10 PCR cycles. Second PCR products were pooled and gel purified from a 2% agarose gel using the MinElute Gel Extraction kit (QIAGEN). Purified product concentration was measured using a Qubit (Thermo Scientific). All products were normalized for each biological sample before combining uniquely barcoded separate biological samples. The pooled products with 10-20% PhiX were sequenced on HiSeq 2500 system (Illumina).

### Generation of artificial antigen expression lentiviral vectors

Plasmids (pCI-neo-sOVA, pCI-neo-mOVA, pCI-neo-cOVA, pcDNA3-OVA) were obtained from Addgene. Different forms of artificial tumor antigen ovalbumin sequence were subcloned into a lentiCRISPR-V2-blast vector via Gibson assembly to generate different Ova-expressing vectors (lentiV2-blast-sOva, lentiV2-blast-mOva, lentiV2-blast-cOva, lentiV2-blast-Ova).

### Generation of artificial tumor antigen Ova-expressing cell lines

4T1, EMT6 and MC38 murine cancer cells were transduced with artificial tumor antigen Ova-expressing lentivirus for 24 hours. After blasticidin selection for 3 days, transduced cancer cells were cultured individually in 10 cm tissue culture plates. One week later, ovalbumin expression levels of transduced cancer cells were identified by immunoblotting.

### *In vivo* CRISPR screening in murine cancer cells

Transduced murine cancer cells were expanded in vitro for 1 week to allow genome editing before being implanted into animals. Cancer cells were either injected into the mammary fat pads of mice or subcutaneously with Matrigel (1:1 dilution). Cancer cells were implanted into both flanks of 10-12 Foxn1^nu/nu^ mice, 10-12 wild-type mice, 10-12 wild-type mice treated with ovalbumin, and 10-12 wild-type mice treated with ovalbumin and PD-1 blockade. Cancer cells transduced with libraries were also grown in vitro at approximately 1000X library coverage for the same time period as the animal experiment. Mice were vaccinated with ovalbumin twice (once a week) 14 days before cancer cell transplantation. Subsequently, mice were treated with 100 μg of rat monoclonal anti-PD-1 (clone: 29F.1A12) on days 9 and 12 via intraperitoneal injection. Mice were euthanized 16–19 days after tumor implantation and tumor genomic DNA was isolated from whole tumor tissue using a DNA extraction protocol (see above). PCR was used to amplify the sgRNA region and sequencing to determine sgRNA abundance was performed on an Illumina HiSeq. Significantly enriched or depleted sgRNAs from any comparison of conditions were identified using the MAGeCK algorithm.

### Mouse validation assays

Ten thousand cancer cells (4T1, MC38) were either injected into the mammary fat pads of mice or subcutaneously with Matrigel (1:1 dilution). Tumors were measured every three days beginning on day 7 after challenge until time of death. Death was defined as the point at which a progressively growing tumor reached 2.0 cm in the longest dimension. Measurements were taken manually by collecting the longest dimension (length) and the longest perpendicular dimension (width). Tumor volume was estimated with the formula: (L × W^2^) / 2. CO_2_ inhalation was used to euthanize mice on the day of euthanasia. Optimal group sizes were determined empirically. Researchers were not blinded to group identity and randomization of animal groups was done when appropriate.

### Cell Viability Assays

Cancer cells were seeded in 96-well plates (500 cells per well for short time proliferation or 100 cells per well for long time proliferation), cultured 4 or 8 days before cell counting, and biologically replicated three times. For cell counting, samples were subjected to Cell Counting Kit 8 (Dojindo) following the manufacturer’s protocol.

### Western blot of protein expression in murine cancer cells

Pellets from 5 × 10^6^ cells were collected and digested by 500 μl RIPA Buffer (Invitrogen). Samples were incubated on ice for at least 15 minutes and centrifuged at 12,000 rpm for 10 minutes at 4 °C, then subjected to BCA analysis (Thermo scientific, Cat# 23228). Approximately 40 μg of total protein from each sample was loaded for western blot analysis.

### Tissue processing and flow cytometry

Tumors for flow cytometry were broken down into smaller fragments, about the size of lentils, then dissociated with 1 mg/ml Collagenase IV for 30 minutes using GentleMacs Octo Dissociator from Miltenyi, and cell suspensions were passed through 70 μm filter twice before staining. Single cancer cells were washed with ice-cold PBS with 2% FBS and stained with antibodies at 4°C for 30 minutes. Cancer cells were then washed and resuspended in ice-cold PBS with 2% FBS for flow cytometry. All data acquisition was done using an LSR II (BD) or FACS Calibur (BD Biosiences) and analyzed using FlowJo software (TreeStar) for statistical computing.

### *In vivo* competition assays

Cancer cells were engineered to express EGFP or mCherry by lentiviral transduction to differentiate populations. Cas9-target sgRNA-transfected cells and Cas9-control sgRNA-transfected cells were mixed and then grown for at least two passages in vitro before implantation into mice. Mixes were analysed by flow cytometry on the day of tumor inoculation. Tumors were harvested and incubated in Collagenase IV for at least 30 minutes. After incubation, cancer cells were passed through 70 μm filters to remove undigested tumors. Single cancer cells were washed with ice-cold PBS with 2% FBS and stained with Near-IR Live/Dead (BD Biosciences) on ice for 30 minutes. Cancer cells were then washed and resuspended in ice-cold PBS with 2% FBS. An LSR II (BD Biosciences) was used to analyze final EGFP/mCherry cancer cell ratios.

### *In vitro* and *in vivo* chemokine measurement

Chemokine expression levels in the culture supernatants were measured using the Mouse Cytokine Array C1000 (Raybiotech). This assay was used to quantify the concentration of chemokines secreted by cancer cells, according to the manufacturer’s instructions. The results were further normalized with protein concentration of tumor cell lysates in the same experiment.

### Generation of CRISPR/Cas9 Knockout cells

Construction of lenti-CRISPR/Cas9 vectors targeting Cop1 was performed following the protocol associated with the backbone vector lentiCRISPR V2 (49535, Addgene). The sgRNA sequences used are listed in the Key resources table. 4T1 and MC38 cells were infected with lentivirus expressing sgRNAs targeting Cop1. After puromycin selection, cells were expanded and collected, and knockout was verified by western blot analyses.

### RNA-seq

Total RNA was isolated and purified from the cells using Isol-RNA Lysis Reagent (Fisher) and treated with DNase I (Fisher). RNA-seq libraries were prepared using the TrueSeq Stranded Total RNA Library Prep Kit (Illumina) and sequenced on an Illumina HiSeq 2500 with 150 base paired end reads.

### Real-time reverse transcription-PCR

RNA was extracted using RNeasy Plus Mini Kit (Qiagen) from 4T1 cells. Then, RNA was reverse transcribed into cDNA using iScript™ cDNA Synthesis Kit (Bio-Rad Laboratories). Approximately 50 ng cDNA from each sample was mixed with gene-specific primers (Supplementary Table 7) and SsoAdvanced™ universal SYBR^®^ Green supermix (Bio-Rad Laboratories) following the manufacturer’s protocol. Reactions were performed on a CFX96 Touch Real-Time PCR Detection System (Bio-Rad Laboratories).

### ATAC-seq

Mouse 4T1 cells were seeded onto 6-well plates for 3 days. Each sample of 1 x 10^5^ cells was trypsinized and resuspended in 50 uL cold ATAC-resuspension buffer (RSB) (10 mM Tris-HCl pH 7.4, 10 mM NaCl, and 3 mM MgCl2 in water) supplemented with 0.1% NP40, 0.1% Tween-20, and 0.01% digitonin. After 3 minute incubation on ice, 1 mL ATAC-RSB containing 0.1% Tween-20 was added, and centrifuged for 10 minutes at maximum speed at 4**°**C. Supernatant was removed and nuclei were resuspended in 50 μL of transposition mix: 2.5 μL transposase (100 nM final), 16.5 μL 1X PBS, 0.5 μL 1% digitonin, 0.5 μL 10% Tween-20, and 5 μL water. Transposition reactions were performed at 37 **°**C for 30 minutes in a thermomixer, while shaking at 1000 rpm. Sequencing libraries were constructed as described. All samples were sequenced using an Illumina HiSeq 2500 with 150 base paired end reads.

## QUANTIFICATION AND STATISTICAL ANALYSIS

### Software used in this study

cutadapt v1.8.1, bowtie2 v2.3.3, samtools v1.9, picard v1.123, MACS2 v2.1.0.20150731, Tophat2 v2.0.11, HT-seq v0.6.1p1, DEseq2 1.22.2, BWA, GATK, MuTect v1.1.4, ROSE v0.1, Cell Ranger v2.0.2, Seurat v2, MAGeCK v0.5.7, BETA, LISA

### CRISPR screen data analysis

CRISPR data were analyzed by MAGeCK and MAGeCK-VISPR essentially as described (Li et al., 2014; Li et al., 2015). Briefly, raw sequencing data are pre-processed by using MAGeCK to obtain the read counts for each sgRNA. Control sgRNAs are used to normalize the data. MAGeCK TEST algorithm is used to compare treatment with control samples to obtain the significantly enriched and depleted sgRNAs and genes. Genes with p value less than 0.001 are candidate hits. The MaGeCKFlute package was used to visualize the data (Wang et al., 2019).

### RNA-seq analysis

Raw RNA-seq reads are aligned to version hg19 of the human genome by using Tophat2 with the default parameters (Kim et al., 2013). Gene counts are quantified by using HT-seq with 44 REFSEQ annotations (Anders et al., 2015; Kim et al., 2013). Differentially expressed genes are identified by using DESeq2 with cutoff of q value < 0.001, ranked by the statistics (Love et al., 2014).

### ATAC-seq analysis

ChiLin pipeline 2.0.0 is used for QC and preprocess of the ATAC-seq (Qin et al., 2016). We use Burrows-Wheeler Aligner (BWA) as a read mapping tool (Li and Durbin, 2009; Qin et al., 2016), and Model-based Analysis of ChIP-Seq (MACS2) as a peak caller (Zhang et al., 2008), with a q-value (FDR) threshold of 0.01. Based on a dynamic Poisson distribution MACS2 can effectively capture local biases in the genome sequence, allowing for more sensitive and robust prediction of binding sites. Unique read for 48 a position for peak calling is used to reduce false positive peaks, statically significant peaks are finally selected by calculated false discovery rate of reported peaks. Deeptools is used for the heatmap plots (Ramírez et al., 2016; Zhang et al., 2008). ATAC-seq Peaks from all study samples were merged to create a union set of sites. Read densities were calculated for each peak for each sample, differential peaks between WT and KO were identified by DEseq2 with adjusted P ≤0.05, |log2fold change| ≥ 1.

### Cop1 degron motif prioritization

#### Cop1 Degron Motif Search

A degron sequence motif for Cop1 was downloaded from the Eukaryotic Linear Motif Database (Gouw et al., 2018), which was represented by the regular expression “[STDE]{1,3}.{0,2}[TSDE].{2,3}VP[STDE]G{0,1}[FLIMVYPA]”. Protein sequences from the mouse and human proteome were downloaded (10/2/2019) from the Swiss-Prot reviewed sequences of the UniProt database (UniProt Consortium, 2019). The Cop1 degron sequence motif was then searched against Swiss-Prot sequences using the python “re” package. This resulted in 1,196 hits (1,067 genes) in mice and 1,328 (1,010 genes) in humans.

### Machine learning prioritization

Not all instances of a sequence motif may be a biologically plausible degron. To further refine plausible candidates, we developed a model to predict the potential of a motif to be a degron. A Random Forest algorithm was trained (number of trees=1,000) on 83 features from the SNVBox database (UniProt Consortium, 2019; Wong et al., 2011) to distinguish previously reported degrons (n=186) (Mészáros et al., 2017) from random other sequences within the same set of proteins (n=186). Features spanned characterization of evolutionary conservation to biophysical features of amino acid residues within a protein. To summarize features across the multiple amino acid residues in a motif, we took the average of each feature. Evaluated using 20-fold cross-validation, performance as measured by the area under the Receiver Operating Characteristic curve (auROC) was 0.81 out of 1.0 (p=2×10^−25^, Mann-Whitney U test).

### Cop1 Degron Motif Filtering

Given the large number of motif hits found in mice, we filtered out those not also seen in humans or which had low degron potential according to machine learning predictions. Of the 1,067 genes with motif hits in mice, 448 showed overlap in humans. After filtering for a high Random Forest score (>0.5 out of 1.0), 117 high-scoring motifs remained. Among the high scoring candidates, numerous were for previously reported Cop1 substrates, such as Ets1, Etv5 and Jun (Marine, 2012).

### Establishing a Tribbles degron motif

Given a prominent hit of a tribbles protein (Trib2) which is known as substrate adaptor of Cop1 (Yokoyama and Nakamura, 2011), we expanded our motif search to find tribbles motifs that may serve a degron recognition purpose for substrate degradation. A previous study has found COP1-TRIB1 binds and degrades the transcription factor CEBPA (Yoshida et al., 2013), with several amino acid residues identified as important for binding (Murphy et al., 2015; Yoshida et al., 2013). We therefore analyzed the multiple sequence alignment of these important residues to generate a plausible degron sequence motif (regular expression: “[IML]…E.[SAT][IL].[IFL]…[IL]”), given the tribbles-mediated degradation of a CEBP family transcription factor is still conserved in drosophila melanogaster (Rørth et al., 2000). The multiple sequence alignment was generated by Clustal Omega using default parameters (https://www.ebi.ac.uk/Tools/msa/clustalo) (Sievers et al., 2011), and visualized using Jalview (Waterhouse et al., 2009). Searching the motif against UniProt sequences found hits for 62 unique genes of which 36 were expressed in the mass spectrometry data.

### Gene signature analysis

#### Cop1 gene expression signature

We created an RNA expression gene signature for Cop1 KO based on the top 500 differentially expressed genes (250 up-regulated, 250 down-regulated). Genes within the signature were weighted by their log2 fold change values reported by DESeq2 in the Cop1 KO vs Rosa26 (control) condition without IFNG treatment. Only mouse genes with a corresponding human gene were used. A Cop1 signature score was computed by a weighted linear combination of Cop1 KO log2 fold changes with normalized expression values from TCGA (see below).

### TCGA expression data

RSEM quantifications (v2) for RNA expression data from The Cancer Genome Atlas (TCGA) was downloaded from the genomic data commons (https://portal.gdc.cancer.gov/). RNA expression was then log2 normalized, followed by subtracting the median expression value for each gene across the cohort.

### Correlation with Immune Cell Infiltration

Immune cell infiltration was inferred from bulk RNA-seq data using the immunedeconv R package (Sturm et al., 2019), which contains estimates based on 6 different methods (CIBERSORT (absolute mode), TIMER, xCell, EPIC, MCP-counter, quanTIseq) (Aran et al., 2017; Becht et al., 2016; Finotello et al., 2019; Li et al., 2016; Newman et al., 2015; Racle et al., 2017). Cop1 signature scores were then analyzed for their correlation with immune cell infiltration estimates, after adjusting for tumor purity (partial spearman correlation, see (Li et al., 2017)). Benjamini-Hochberg correction was then applied across all p-values and a correlation was deemed significant at a FDR <0.05.

### Cop1 M2 Macrophage-specific Signature

Notably, multiple macrophage infiltration estimates were correlated with the Cop1 KO gene expression signature across multiple cancer types in TCGA. Furthermore, lower estimates for M2 macrophages were associated with better survival in basal-like breast cancer. We therefore sought to examine whether genes differentially expressed by Cop1 KO could also predict the imbalance of M2 macrophages to M1 macrophages in basal-like breast cancers from TCGA. Using LASSO (scikit learn python package), we generated a consensus prediction for two different measures of M2 macrophage imbalance: 1) CIBERSORT estimates by computing a score of M2 macrophages minus M1 macrophages; 2) a previously reported signature of Tumor Associated Macrophage M2 (TAM M2) (Jiang et al., 2018). Only expressions of genes that were both differentially expressed upon Cop1 KO and a cytokine/receptor (see below) were used as features. The regularization parameter lambda in LASSO was determined using 10-fold cross-validation. Performance was then measured on 30% held-out data that was never used for training. The final regression coefficients for genes with non-zero values in both models were then averaged.

### Cytokine and surface receptor/ligand genes

To analyze genes that may impact the tumor-immune microenvironment, we curated a set of cytokine and surface receptor related genes: “Cytokine-Cytokine Receptor Interaction” pathway from KEGG database and surface receptor/ligand genes as reported by Ramilowski et al (Kanehisa and Goto, 2000; Ramilowski et al., 2015).

## Notes

### Competing Interest Statement

The authors have declared no competing interest.

