## Supplemental figures for "*In Vivo* CRISPR Screens Identify E3 Ligase Cop1 as a Modulator of Macrophage Infiltration and Cancer Immunotherapy Target"

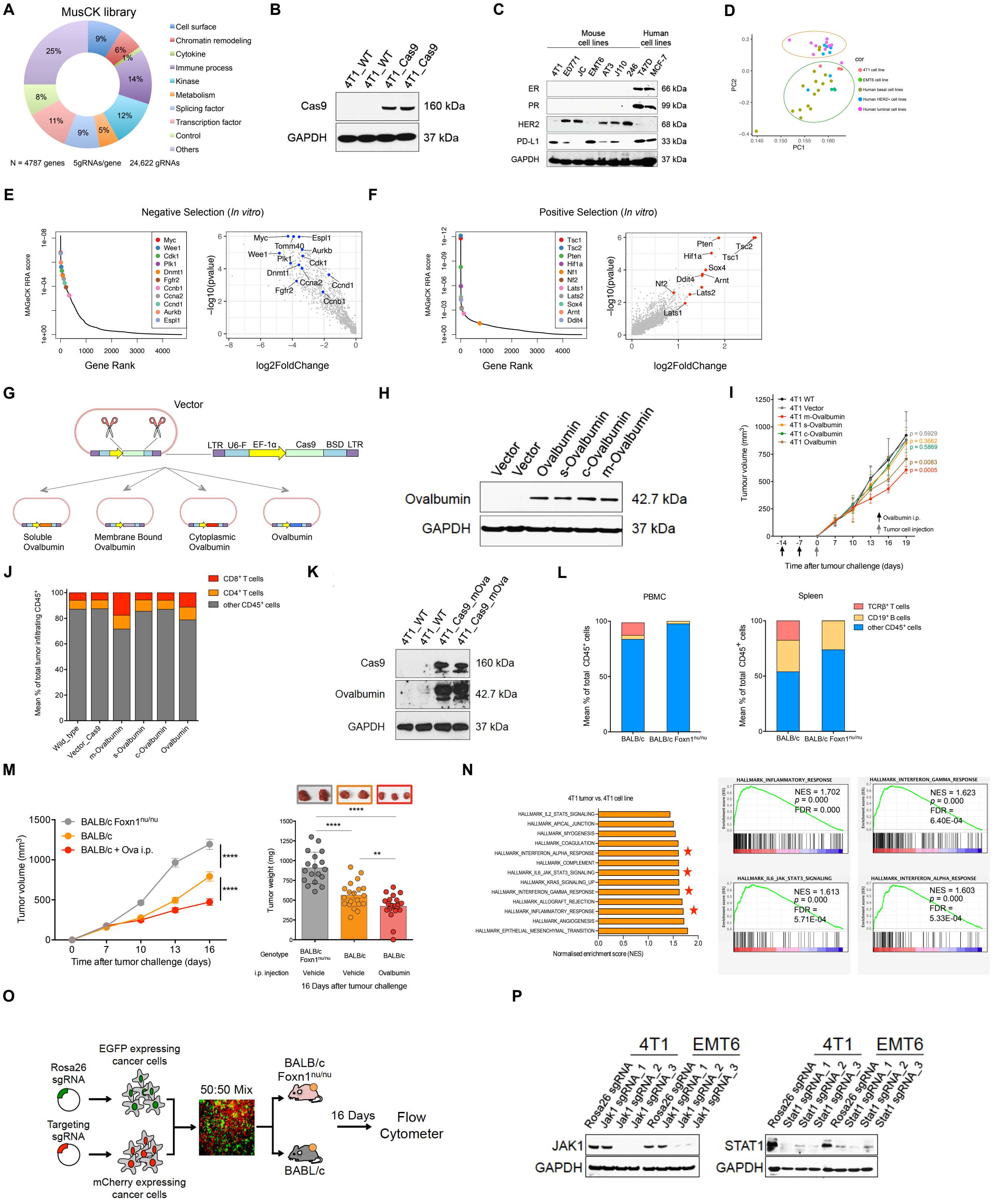


**Figure S1 related to Figure 1**

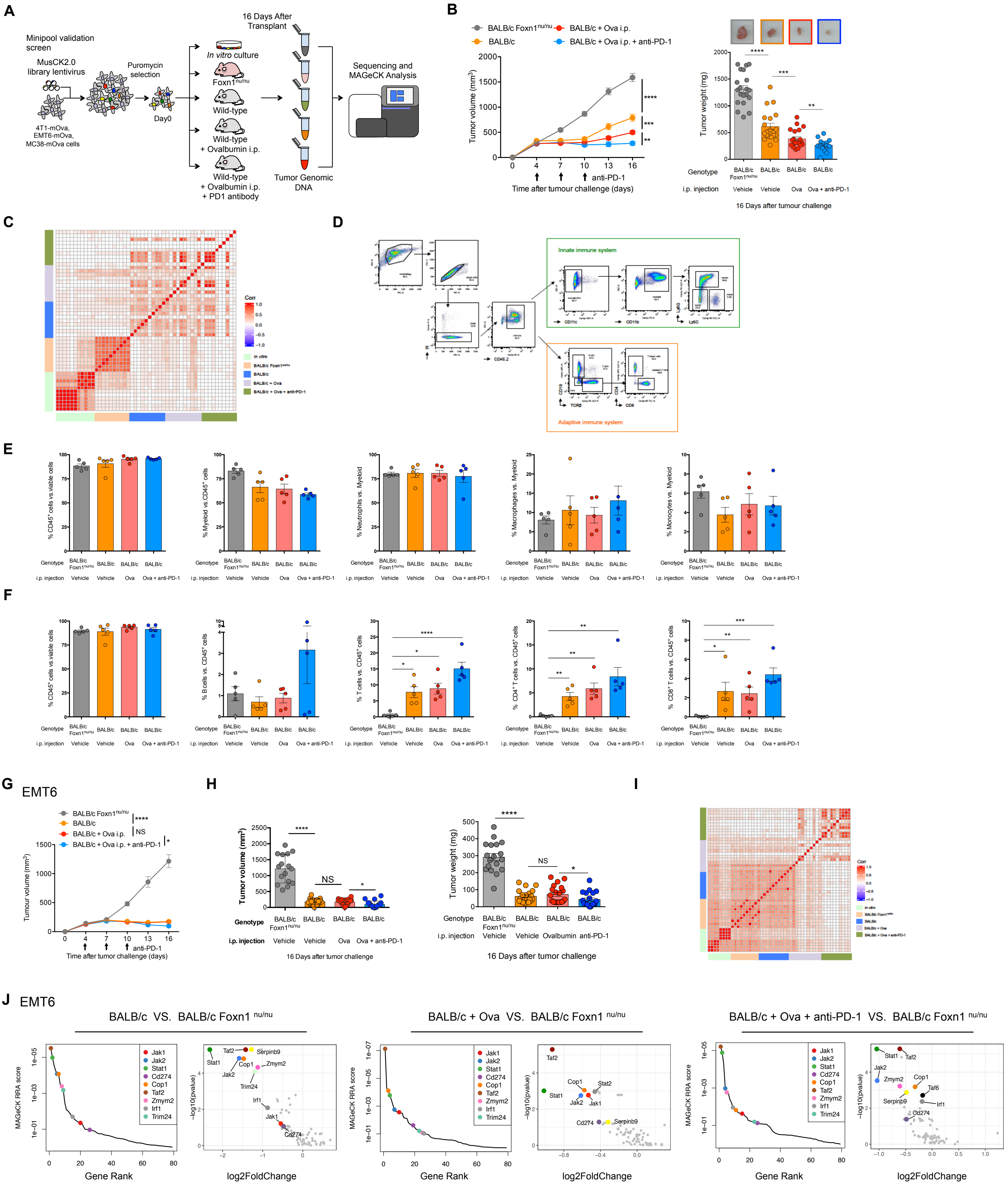


**Figure S2** related to Figure 2

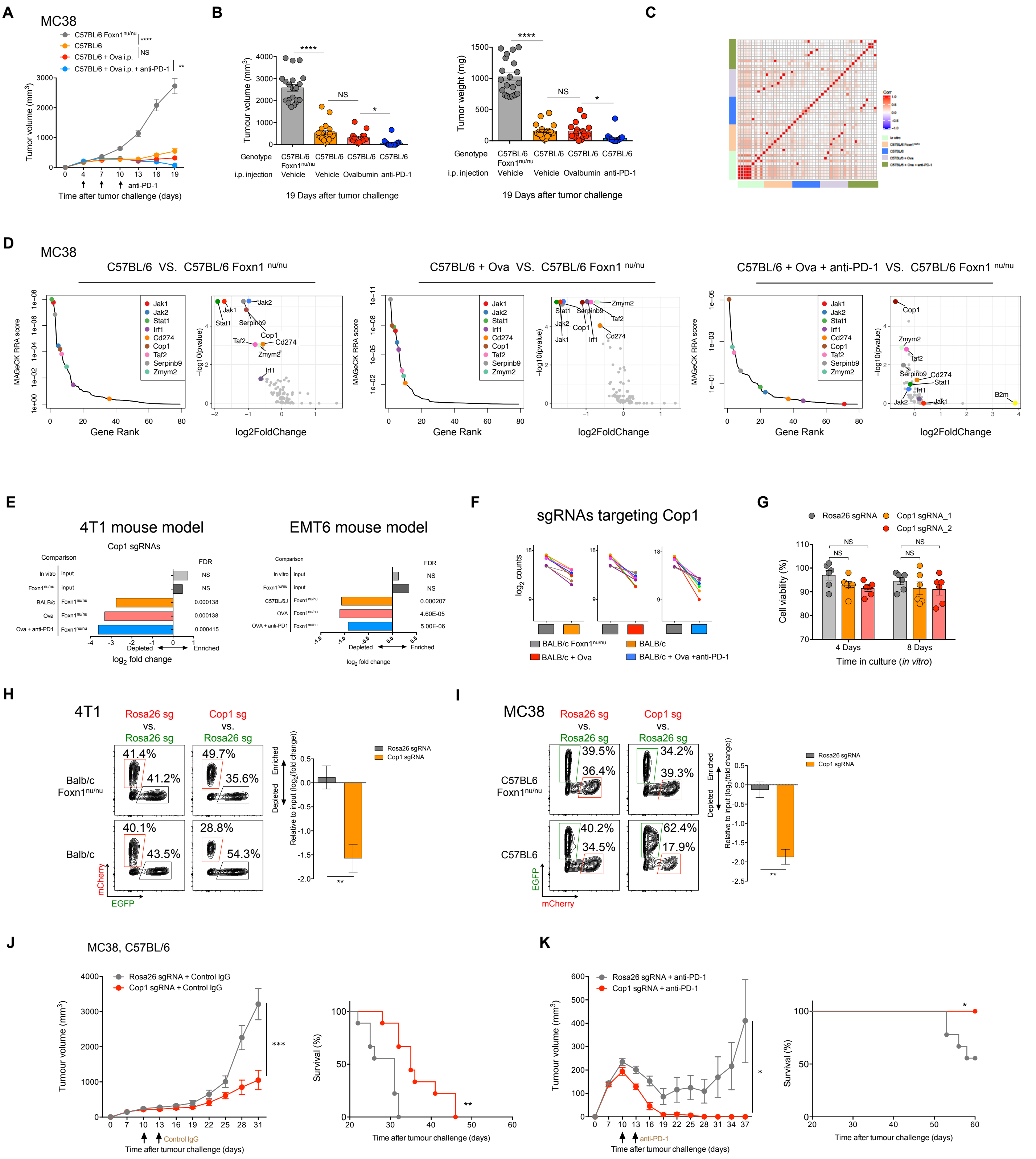


**Figure S3** related to Figure 2

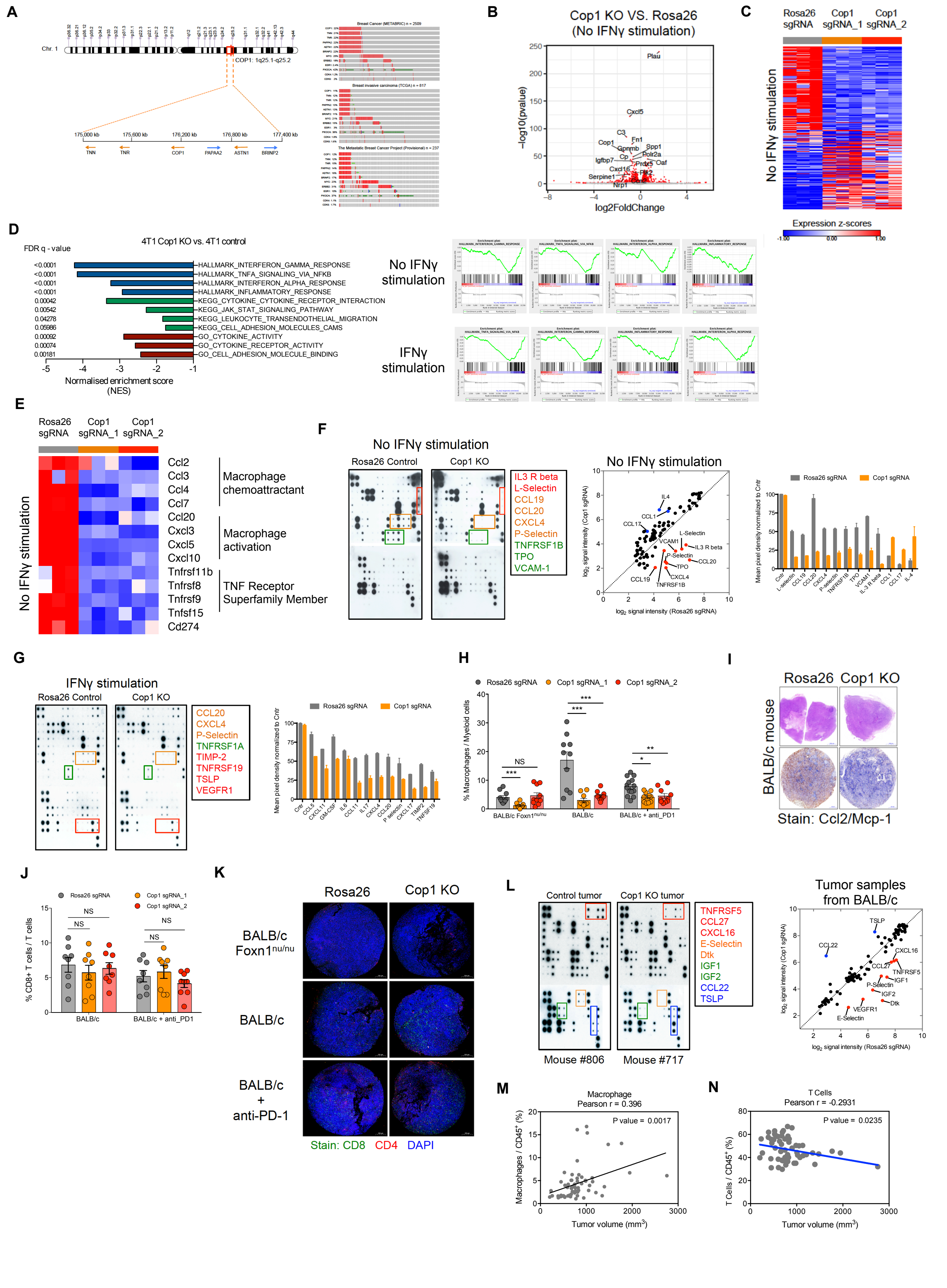


**Figure S4** related to Figure 3

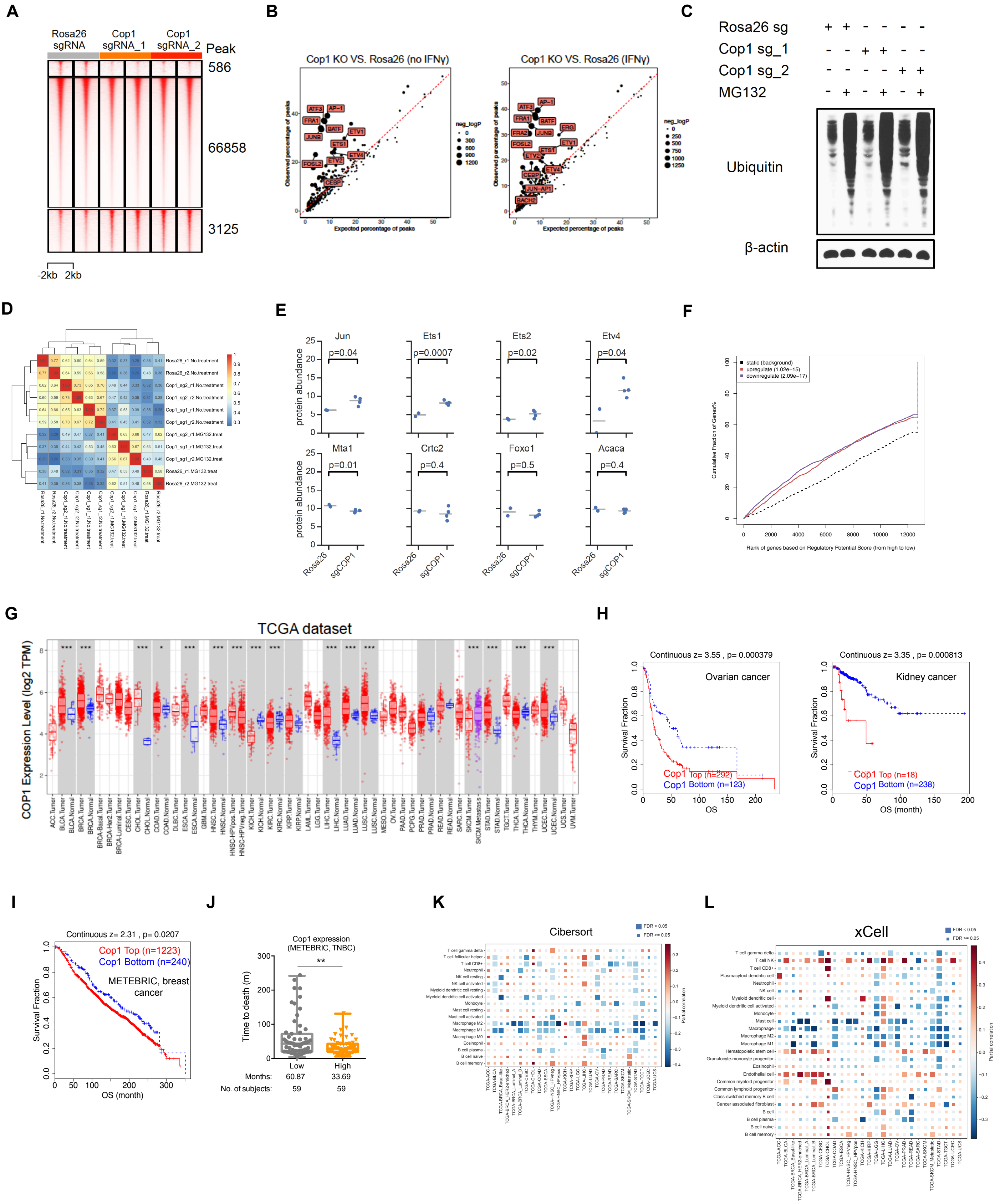


Figure S5 related to Figures 4 and 5

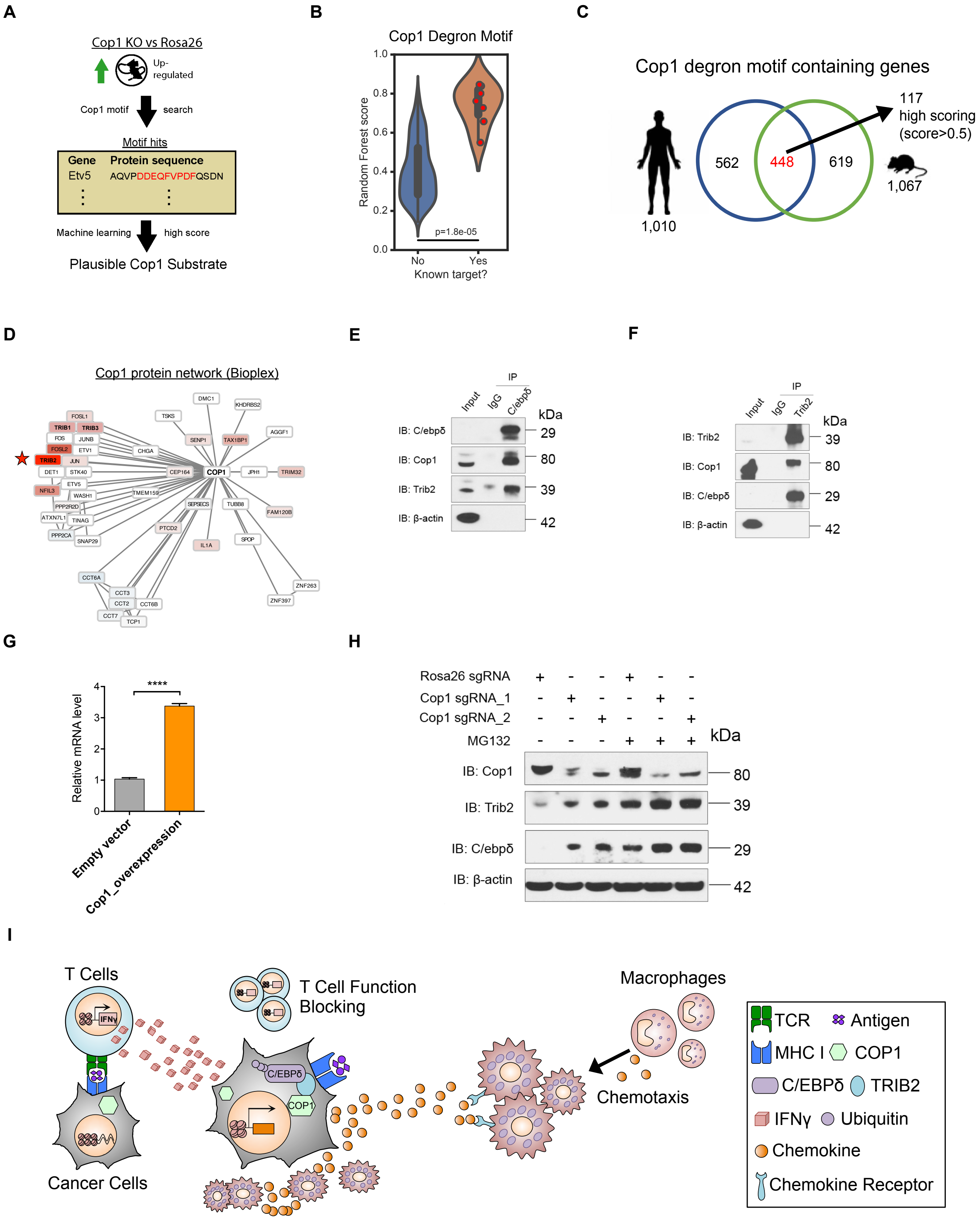


**Figure S6** related to Figure 6
